# Cooperative binding of TCR and CD4 to pMHC enhances TCR sensitivity

**DOI:** 10.1101/2021.11.22.469547

**Authors:** Muaz Nik Rushdi, Victor Pan, Kaitao Li, Hyun-Kyu Choi, Stefano Travaglino, Jinsung Hong, Fletcher Griffitts, Pragati Agnihotri, Roy A. Mariuzza, Yonggang Ke, Cheng Zhu

**Affiliations:** Wallace H. Coulter Department of Biomedical Engineering, Georgia Institute of Technology and Emory University, Atlanta, GA, USA; Parker H. Petit Institute for Bioengineering and Biosciences, Georgia Institute of Technology, Atlanta, GA, USA; Georgia W. Woodruff School of Mechanical Engineering, Georgia Institute of Technology, Atlanta, GA, USA; W. M. Keck Laboratory for Structural Biology, Institute for Bioscience and Biotechnology Research, University of Maryland, Rockville, MD, USA; Department of Cell Biology and Molecular Genetics, University of Maryland, College Park, MD, USA; Medical Scientist Training Program, Emory University School of Medicine, Atlanta, GA, USA

## Abstract

Antigen recognition of CD4^+^ T cells by the T cell receptor (TCR) can be greatly enhanced by the coreceptor CD4^1–7^. Yet, understanding of the molecular mechanism is hindered by the ultra-low affinity of CD4 binding to class-II peptide-major histocompatibility complexes (pMHC)^1,7–10^. Using two-dimensional (2D) mechanical-based assays, we determined a CD4–pMHC interaction to have 3-4 logs lower affinity than cognate TCR–pMHC interactions^8^, and to be susceptible to increased dissociation by forces (slip bond)^5,8,11^. In contrast, CD4 binds TCR-prebound pMHC at 3-6 logs higher affinity, forming TCR–pMHC–CD4 trimolecular bonds that are prolonged by force (catch bond)^5,8,11^ and modulated by protein mobility on the cell membrane, indicating profound TCR-CD4 cooperativity. Consistent with a tri-crystal structure^12^, using DNA origami as a molecular ruler to titrate spacing between TCR and CD4 indicates that 7-nm proximity optimizes trimolecular bond formation with pMHC. Our results reveal how CD4 augments TCR antigen recognition.

## Main

CD4^+^ T cells play vital and versatile roles in the adaptive immunity by differentiating into lineages of various functions upon activation by antigen. Antigen recognition is achieved by the T cell receptor (TCR) interacting with its cognate peptide bound to major histocompatibility complex (MHC, or pMHC when including the peptide) expressed on an antigen-presenting cell (APC) or target cell. TCR–pMHC binding triggers phosphorylation of CD3 immunoreceptor tyrosine-based activation motifs (ITAMs), leading to a signaling cascade that results in T cell activation and effector function^13^. Expressed on the surface of their respective T cell subsets, CD8 and CD4 coreceptors are thought to facilitate the initiation of the intracellular signaling cascade by stabilizing TCR–pMHC binding and delivering its associated Lck to TCR^14,15^. However, unlike CD8, CD4’s extremely low affinity for MHC hinders our understanding of these proposed mechanisms that require CD4 binding to pMHC^9,16^. Indeed, surface plasmon resonance (SPR) was unable to measure CD4–MHC interactions in three-dimensions (3D)^7,17^, except for one report of a ~200 μM *K*_d_^10^ that was not reproduced^7^. Although affinity measurements were successfully attained using B cells and a supported lipid bilayer reconstituted with CD4^7^, it required additional CD2–CD58 interactions to stabilize the 2D interface. The estimated 2D *K*_d_ of ~5,000 μm^−2^ is >1-log higher than the *K*_d_ of TCR–pMHC interactions^18^ or the CD4 density of T cell surface^8^, neither of which predicts CD4–MHC interaction *in vivo*.

Also due to the extremely low CD4–MHC binding affinity, the ectodomain crystal structure of TCR–pMHC–CD4 tri-molecular complex was not available until a gain-of-function CD4 mutant with substantially higher affinity toward MHC was engineered^12^. Fascinatingly, the rigid coreceptor maintains a membrane proximal spacing of ~7 nm apart from the TCR while binding to the MHC, providing an adequate gap for the missing CD3 proteins to amalgamate around the TCR^17,19^. The co-crystal structure suggests that the membrane proximal positioning of TCR and CD4 ectodomains is energetically favorable to form a trimolecular complex with pMHC. Due to the differential affinities to pMHC, it is unlikely driven by the CD4 binding, but instead needs the TCR binding to pMHC to initiate the trimolecular complex formation. The above reasoning provides a rationale of our hypothesis that CD4 augmentation comes into play via its synergy with TCR, a mechanism similar to that of TCR-CD8 cooperation^3–5,20^. However, such cooperativity remains elusive with conflicting observations from previous studies. Our previous study using a mechanical 2D assay showed no effect of murine CD4 on murine TCR–pMHC binding under force-free conditions^8^. Studies of others using Forster Resonance Energy Transfer (FRET)^18^ and pMHC tetramer staining techniques^21^ failed to detect CD4 enhancement, whereas another study using Fluorescence Recovery After Photobleaching (FRAP) observed that TCR and CD4 reciprocally enhance each other’s binding to pMHC^22^. Most importantly, CD4’s ability to bind pMHC is still required for TCR-mediated IL-2 production even when its association with Lck is abolished^22^.

Combining our 2D kinetic assays and mathematical modeling, we analyzed the bimolecular CD4–MHC interaction and TCR-CD4 cooperativity using both purified protein ectodomains and cell systems. We identified enormous cooperative binding of CD4 to TCR-prebound pMHC, which has 3-6 logs higher affinity than binding to free pMHC. This cooperativity manifests substantially greater number and longer-lived bonds of pMHC with both TCR and CD4 than the sum of separate pMHC bonds with TCR or CD4, which results from the emergence of a dominating TCR–pMHC–CD4 trimolecular bond species that is dynamically regulated by mechanical force applied on the molecular complex. By presenting TCR and CD4 on different surfaces to modulate molecular diffusion or DNA origami to modulate molecular spacing, we showed that the TCR-CD4 cooperativity requires molecular mobility to allow for their colocalization within a 7-nm proximity, consistent with that reported in TCR–pMHC–CD4 tri-crystal structure. Our results explain how CD4 could efficiently augment TCR antigen recognition via cooperative binding to TCR-prebound pMHC, thereby boosting TCR sensitivity. They also underline the importance of physical factors in regulating T-cell antigen recognition, which may be imposed on molecular organization at the cell membrane via signaling dependent and/or independent mechanisms.

## Results

### CD4–MHC binding displays low 2D affinity

We used our adhesion frequency assay^23,24^ (Fig. 1a), which analyzes cross-junctional interactions upon initial brief cell-cell encounters, to quantify the 2D binding between HLA-DR1 presenting a melanoma antigenic peptide derived from the glycolytic enzyme triosephosphate isomerase (TPI:HLA-DR1, referred hereafter as pMHC, unless specified)^25^ and its cognate E8 TCR, wild-type (WT) CD4, or a high-affinity mutant (MT) CD4 generated via directed evolution^17^. Ectodomains of each binding pair were respectively coated on two opposing RBCs, which were brought into contact and then separated to detect adhesion by RBC elongation. Results were presented as adhesion frequency (*P*_a_) of many repeated contact-separation cycles, which was higher between RBCs functionalized with designated molecules at appropriate densities than between blank ones at the same contact time *t*_c_, indicating specific interactions (Fig. 1b). *P*_a_ increased initially and plateaued as the probability of bond formation was balanced by that of dissociation with prolonged *t*_c_, resulting in a steady-state (Fig. 1c). We evaluated the effective 2D affinity (*A*_c_*K*_a_) and off-rate (*k*_−1_) by fitting the *P*_a_ vs *t*_c_ curve (Fig. 1c) or the log-transformed and molecular density-normalized counterparts (Fig. 1d) to a published model^23,24^ (Table 1). The log transformation converts *P*_a_ to the average number of bonds per contact, <*n*> = - ln(1 – *P*_a_), which, after normalization by densities of the receptor (*m*_r_) and ligand (*m*_l_) collapses three curves measured at different *m*_r_×*m*_l_ values (indicated by different symbols in Fig. 1d) and approaches to *A*_c_*K*_a_ at large *t*_c_. This model was well supported as the steady-state <*n*> increased linearly with *m*_r_×*m*_l_ (Supplementary Fig. 1a-c) and collapsed into a single plateau upon normalization by *m*_r_×*m*_l_ (Fig. 1e), suggesting the lack of multimeric binding and supporting the reliability of our measurements^23,24^. The effective 2D affinity of E8 TCR for pMHC (*A*_c_*K*_a,TCR_ = 7.70 ± 0.40 × 10^−4^ μm^4^) is comparable to, but off-rate (*k*_−1,TCR_ = 0.48 ± 0.07 s^−1^) is slower than, other TCR–pMHC-II interactions^8^. Remarkably, the effective 2D affinity of CD4–MHC-II binding (*A*_c_*K*_a,CD4_ = 4.35 ± 0.49 × 10^−7^ μm^4^) is >3 logs lower and among the lowest values recorded^26^ (Fig. 1e). Consistent with its 3D SPR measurement^17^, the affinity of MT CD4 for MHC (2.48 ±0.15 × 10^−3^ μm^4^) is 3-4 logs higher than that of WT CD4 (Fig. 1e, Table 1).

**Figure 1.**
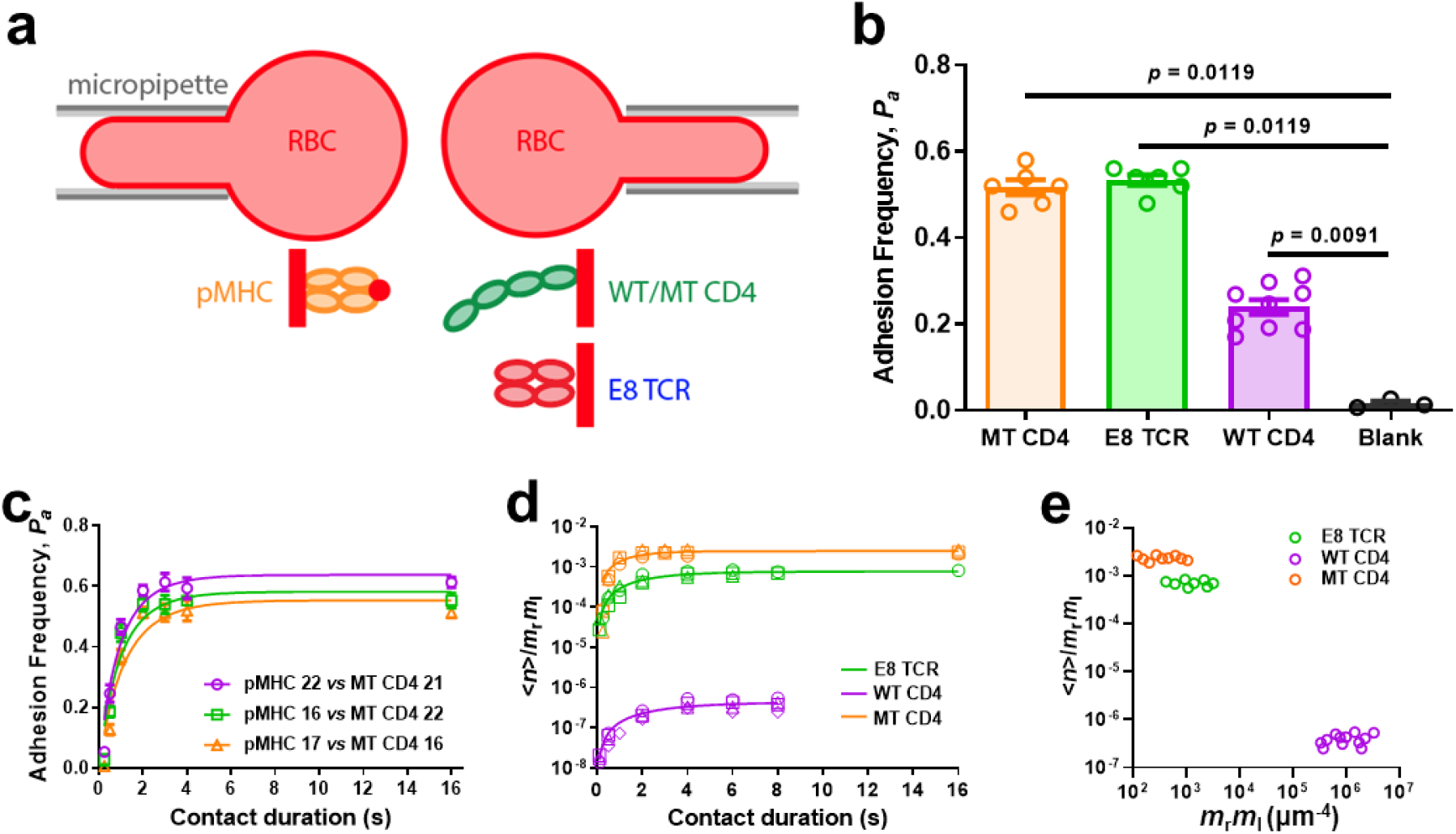
2D kinetic analysis of TPI:HLA-DR1 interaction with E8 TCR or CD4. **a,** Schematics of micropipette binding frequency assay (*top*) and molecular coating (*bottom).* A pMHC-coated RBC (*left*) was repeatedly brought into contact with an opposing RBC coated with E8 TCR, or WT/MT CD4 (right) and separated to detect bond formation across the contact interface. **b,** Specificity is shown by the significant higher (p-value indicated, by Mann-Whitney test) of steady-state (contact time *t*_c_ = 4 s) adhesion frequencies (*P*_a_, mean ± SEM (n = 6, 6, 9, 3 cell pairs) of indicated points each determined from 50 cycles of contact) between pMHC-coated RBCs and RBCs coated with E8 TCR, WT CD4, or MT CD4, than that without ligand. **c,** Mean ± SEM (n = 3 cell pairs per data point) *P*_a_ vs *t*_c_ curves for pMHC interacting with MT CD4 at various coating densities as indicated by the number in legend. **d,** The normalized bond number <*n*>/*m*_r_*m*_l_ vs *t*_c_ curve for pMHC interaction with E8 TCR (green), WT CD4 (purple), or MT CD4 (orange). Different symbols represent independent experiments using different molecular densities that are shown in Supplementary Table 1. **e,** Normalized bond number at steady-state calculated from **d** was plotted vs receptor and ligand densities.

**Table 1.**
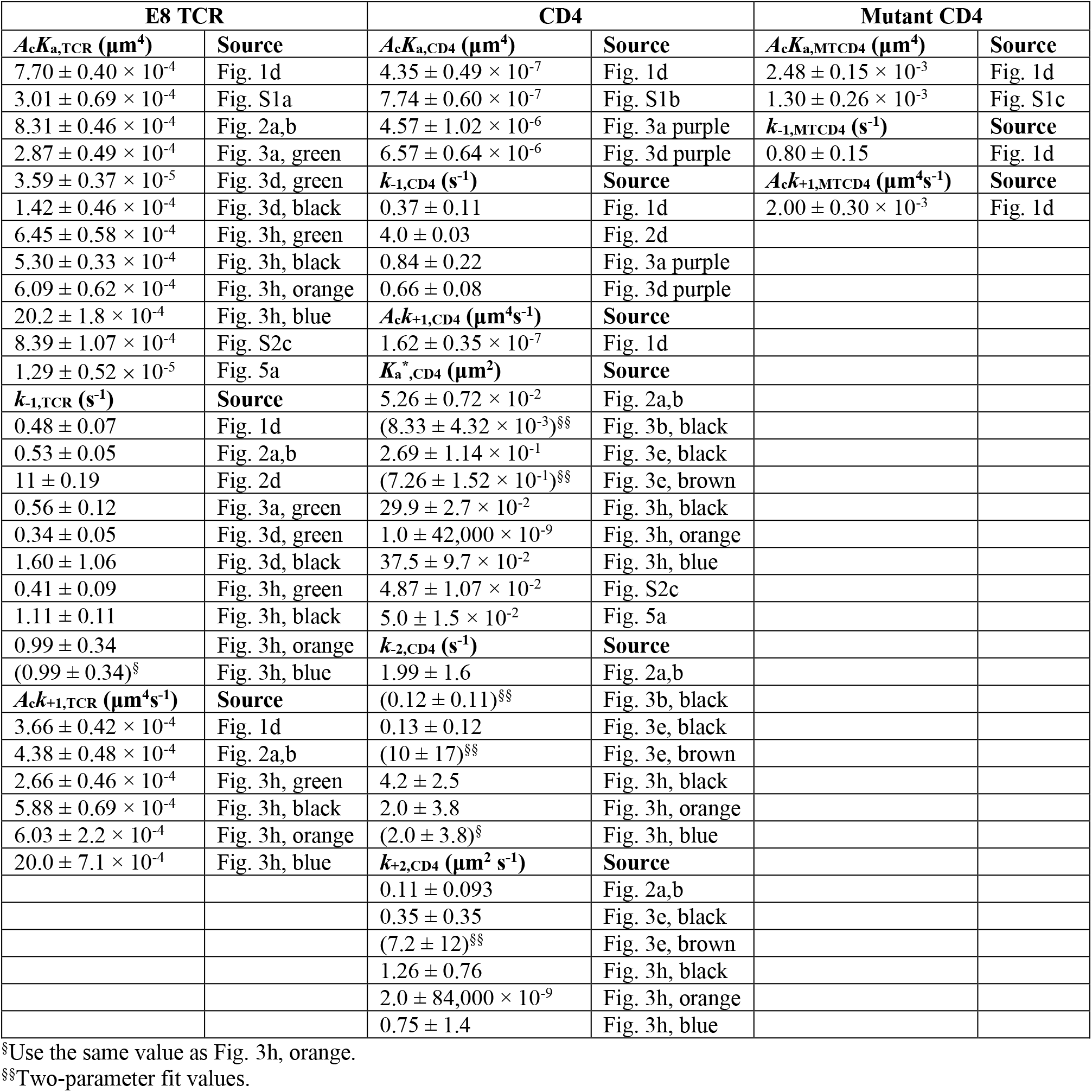
Kinetic parameters of the purified protein system and the measurement methods.

### TCR and CD4 cooperatively bind to pMHC

To explain how CD4–MHC binding could disproportionally recruit Lck to TCR^7^, we tested the hypothesis that TCR and CD4 bind pMHC synergistically as observed for TCR and CD8^3–5^. E8 TCR and CD4 were co-functionalized on RBCs at different density ratios (*m*_TCR_/*m*_CD4_) while keeping their sum constant and tested against RBCs bearing pMHC with the same density (*m*_pMHC_). We first analyzed the *P*_a_ vs *t*_c_ curves so measured (Fig. 2a) using a dual-species concurrent and independent binding model^27–29^, opposite to our synergistic binding hypothesis. Without synergy, this model predicts that the average whole number of bonds <*n*>_W_ would be equal to the sum of its parts, i.e., the average numbers of TCR–pMHC bonds, <*n*>_TCR_, plus the average numbers of CD4–MHC bonds, <*n*>_CD4_. Given the low CD4 density, which was far from enough to make up its low affinity, the measured <*n*>_W_ should have yielded <*n*>_TCR_ as its only detectable component and normalizing it by *m*_TCR_ and *m*_pMHC_ should have collapsed the entire family of curves as did Fig. 1d. Contrary to this prediction based on the dual-species concurrent and independent binding model^27–29^, the <*n*>_W_/(*m*_TCR_×*m*_pMHC_) vs *t*_c_ curve upward shifted as *m*_CD4_ increased (Fig. 2b and Supplementary Fig. 2a), falsifying the independent binding assumption^27–29^. Instead of increasing linearly as in Supplementary Fig. 1, the steady-state <*n*>W increased nonlinearly with increasing *m*_TCR_×*m*_pMHC_ (Supplementary Fig. 2b). Instead of being a constant as Fig. 1e, the TCR- and pMHC-density normalized whole bond number, <*n*>_W_/*m*_TCR_×*m*_pMHC_, increased linearly with increasing CD4 density *m*_CD4_ (Supplementary Fig. 2c). These results demonstrate that TCR and CD4 cooperatively bind to pMHC as we hypothesized, manifesting higher number of whole bonds than the sum of TCR–pMHC and CD4–pMHC bonds as parts.

**Figure 2.**
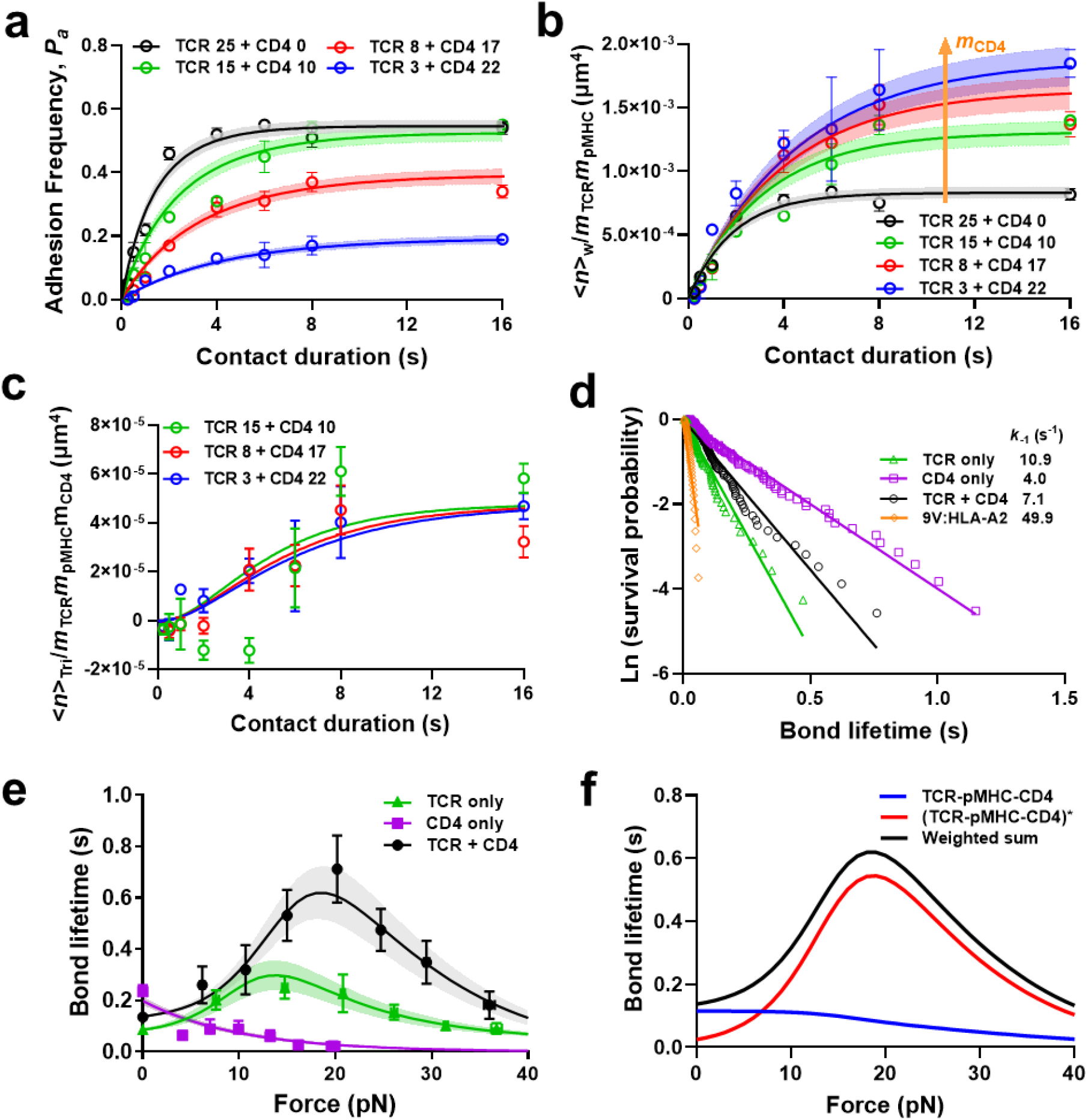
Ectodomains of TCR and CD4 bind cooperatively to pMHC. **a,** *P*_a_ vs *t*_c_ data and their model fits of pMHC-coated RBCs (38 μm^−2^) binding to RBCs coated with TCR alone, or a mixture of TCR and CD4 at densities (# of molecules/μm^2^) indicated by the number in legend. **b,** Normalized whole bond number <*n*>_W_/*m*_TCR_*m*_pMHC_ vs *t*_c_ curves and their respective model fits for interactions in **a**. Arrow indicates decreasing density of TCR and increasing density of CD4. **c,** Normalized trimolecular bond number <*n*>_tri_/*m*_TCR_*m*_pMHC_*m*_CD4_ vs *t*_c_ curves calculated from **b** by subtracting the black data from colored data followed by further normalization by *m*_CD4_. **d**, Survival probabilities plotted as ln(# of events with lifetime greater than *t*) vs bond lifetime for pMHC-coated beads interacting with beads coated with TCR, CD4, or a 1:1 mixture of TCR and CD4 measured by BFP thermal fluctuation at zero force. Each data set (points) was fitted by a straight line with the negative slope representing off-rate for TCR (*k*_−1,TCR_, n = 71) or CD4 (*k*_−1,CD4_, n = 92), or apparent off-rate for the mixed TCR and CD4 (n = 97). As a negative control, class-I MHC 9V:HLA-A2 was used instead of TPI:HLA-DR1 to measure non-specific binding (n = 42). **e,** Mean ± SEM bond lifetime vs force data (points), measured by BFP force-clamp experiment using beads bearing TPI:HLA-DR1 interacting with E8 TCR (n = 230), CD4 (n = 355), or a 1:1 mixture of E8 TCR and CD4 (n = 424) coated on beads, are compared with theoretical curves (mean ± SE) predicted using the model and best-fit parameters in Supplementary Fig. 3. **f**, Model predicted trimolecular bond lifetime vs force plots for strong (red) and weak (blue) states as well as their sum (black). Data (points) in **a** and **b** are presented as mean ± SEM of two independent experiments for each curve. Their model fits (mean ± SE) are calculated using Eqs. 1 and 2 based on the best-fit parameters listed in Table 1.

From the difference between the whole and the sum of parts, we identified a new bond species – the emerging TCR–pMHC–CD4 trimolecular bonds whose average number is <*n*>_tri_ = Δ<*n*> = <*n*>_W_ – (<*n*>_TCR_ + <*n*>_CD4_). We developed a kinetics model in which CD4 binding of free pMHC is ignored due to its negligible affinity compared to TCR binding, but CD4 is allowed to bind TCR-prebound pMHC with a higher affinity 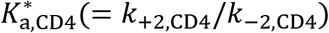. This model predicts (see Methods):

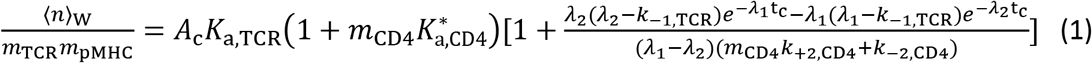

where *λ*_1_ and *λ*_2_ are positive functions of off-rate of TCR–pMHC bond (*k*_−1,TCR_), on- and off-rate of CD4 binding to (*k*_+2,CD4_) and dissociating from (*k*_−2,CD4_) TCR-prebound pMHC, and CD4 density (*m*_CD4_):

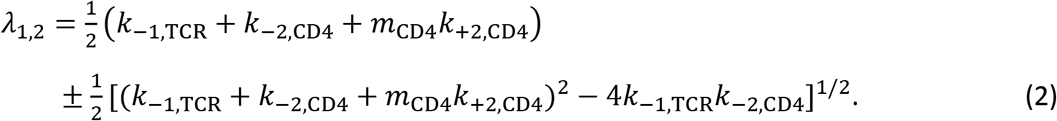

The steady-state solution as *t*_c_ → ∞ is

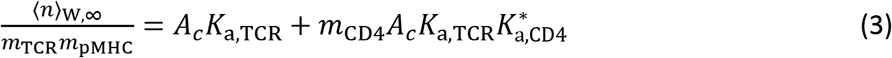

The second term on the right-hand side of Eq. 3 can be identified as the steady-state average number of trimolecular bonds <*n*>_tri_ per densities of TCR and pMHC. Both the original (Fig. 2a) and normalized (Fig. 2b) data were well fitted by Eqs. 1&2. Fitting returned values of *A*_c_*K*_a,TCR_ = 8.31 ± 0.46 × 10^−4^ μm^4^ and *k*_−1,TCR_ = 0.53 ± 0.05 × s^−1^ for the TCR–pMHC interaction from the dual receptor species experiment in the presence of CD4, which agree well with the previous single receptor species experiment in the absence of CD4 (Table 1). Remarkably, the value of 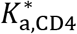 is 0.0526 ± 0.0072 μm^2^, >250 times higher than the previous value obtained using CD4-reconstituted supported lipid bilayers^7^ and >350,000 times higher than the CD4 affinity for free pMHC calculated by dividing the *A*_c_*K*_a,CD4_ value determined from Fig. 1d by the ~3 μm^2^ contact area *A*_c_ estimated from the side-view microscopic image.

To visualize the emergence of TCR–pMHC–CD4 trimolecular bonds, we transformed the data according to Eq. 3: subtracting the black data set (*m*_CD4_ = 0) from the colored data sets (*m*_CD4_ > 0) and normalizing them by their corresponding CD4 density. This nearly collapses the three colored data sets into the average number of trimolecular bonds per unit densities of TCR, pMHC, and CD4. This <*n*>_tri_/(*m*_TCR_×*m*_pMHC_×*m*_CD4_) data vs contact time *t*_c_ plot compares well with our model prediction using the best-fit kinetic parameters obtained earlier, showing how the degree of cooperativity increase in time until reaching the steady-state value 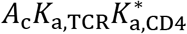 (Fig. 2c). Thus, the steady-state trimolecular bond number per unit density of TCR, CD4, and pMHC is equal to the product of the effective 2D affinity of the TCR–pMHC bond and the 2D affinity of CD4 for TCR-prebound pMHC, an intuitive result that make very good sense.

### TCR–pMHC–CD4 trimolecular bond shows a much greater catch than TCR–pMHC bimolecular bond

To examine whether the trimolecular bond can better sustain mechanical force, we used the biomembrane force probe (BFP) thermal fluctuation^24,30^ and force-clamp^5,8,11^ assays to measure respective force-free and force-dependent lifetimes of single bonds between glass beads bearing pMHC and targets presenting ectodomains of E8 TCR, CD4, or both mixed at a 1:1 concentration ratio. The thermal fluctuation assay yielded fairly linear survival probability vs bond lifetime semi-log plots for dissociation of single-receptor species (TCR or CD4) and dual-receptor species (TCR and CD4) from TPI:HLA-DR1, which were much longer lived than the negative control 9V:HLA-A2 (Fig. 2d), conforming binding specificity of TCR and/or CD4.

The force-clamp assay found the CD4–MHC bond lifetimes to decrease with increasing force, displaying a slip-only bond behavior^5,8,11,31^ (Fig. 2e). The TCR–pMHC interaction, in contrast, behaved as a catch-slip bond where the lifetime increased, reached a peak, and decreased in a biphasic fashion as force increased^5,8,11,31^ (Fig. 2e). Remarkably, the interactions between pMHC probes and targets co-presenting TCR and CD4 exhibited a much more pronounced catch-slip bond (Fig. 2e). Since lifetimes were measured at infrequent binding (<20%) to ensure >89% single-bond probability, the “whole” lifetime ensemble included subpopulations of single TCR–pMHC, CD4–MHC, and TCR–pMHC–CD4 bond lifetimes, where the whole lifetime represented a weighted sum of all three contributions. Average lifetimes of whole bonds were much longer than the weighted sum of two parts – the average lifetimes of TCR–pMHC and CD4–MHC bonds – regardless of how their relative weights were adjusted (Fig. 2e), as it cannot be longer than the longer component lifetime of the two. This demonstrates the existence of a more “catchy” and longer-lived trimolecular bond species.

We used a single-state, single-pathway dissociation model for the slip-only bonds of CD4–MHC interaction and two two-state, two-pathway dissociation models for the catch-slip bonds of the respective TCR–pMHC and TCR–pMHC–CD4 interactions (Supplementary Fig. 3a) to globally fit the measured lifetime vs force scattergrams (Supplementary Fig. 3b). The models fit the mean ± SE bond lifetime vs force data well (Fig. 2e) and predict the probability and fraction of strong state of TCR–pMHC and TCR–pMHC–CD4 bonds (Supplementary Fig. 3e-h) as well as the average bond lifetime vs force curves of the strong state, weak state, and their sum (Fig. 2f), which reveals how the TCR–pMHC–CD4 bond is activated to a strong state by force to form a cooperative catch bond.

### TCR-CD4 cooperativity on cells

We performed two sets of experiments to confirm that the TCR-CD4 cooperativity observed using purified ectodomain proteins was operative when these proteins were expressed on cells. First, we analyzed binding of beads coated with E8 TCR, CD4, or both to THP-1 cells pulsed with TPI peptide after stimulation with IFN-γ to upregulate HLA-DR1 expression^25^. Analyses of adhesion frequency *P*_a_ (Fig. 3a) and of the corresponding normalized bond number <*n*>_W_/*m*_Receptor_×*m*_pMHC_ (Fig. 3b) mediated by the TCR–pMHC (green) and CD4–MHC (purple) bimolecular interactions yielded *A*_c_*K*_a,TCR_ = 2.87 ± 0.49 × 10^−4^ μm^4^ and *A*_c_*K*_a,CD4_ = 4.57 ±1.02 × 10^−6^ μm^4^. The former value is a comparable to, and the latter value is a log higher, than the corresponding values of TCR and CD4, respectively, for the purified MHC.

**Figure 3.**
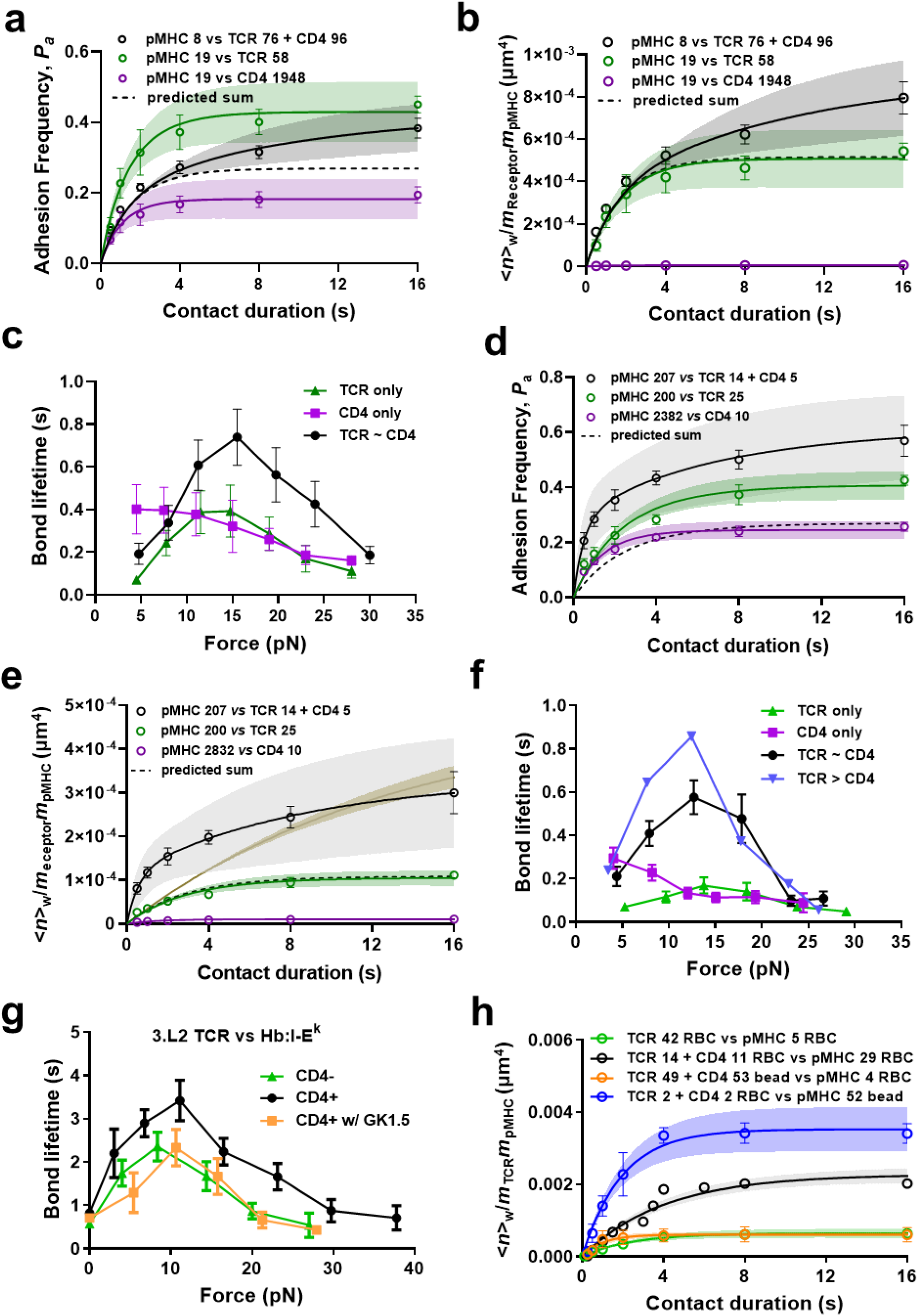
Cellular TCR and CD4 bind cooperatively to pMHC. **a,** *P*_a_ vs *t*_c_ curves (Mean ± SEM of 3 or more cells per data point) and their model fits of TPI-pulsed IFN-γ treated THP-1 cells binding to beads presenting TCR, CD4, or both at densities (μm^−2^) indicated in the legends. The dashed curve represents predicted sum of two concurrent and independent TCR–pMHC and CD4–MHC bimolecular interactions, which used the kinetic parameters of the two bimolecular interactions with molecular densities matched those used in the experiment measuring total interactions. Best-fit *P*_a_ ± SE (curves) was calculated from best-fit <*n*> ± SE in **b**. p = 0.0563, 0.0159, 0.0631, 0.1200, 0.0502, and 0.0405 for comparing measured *P*_a_ of whole interactions to predicted sum at *t*_c_ = 0.5, 1, 2, 4, 8, and 16 s. **b,** Normalized whole bond number <*n*>_W_/*m*_Receptor_*m*_pMHC_ vs *t*_c_ data (points) and their respective model fits (curves) for interactions in **a**. Predications were made after normalizing the densities in different curves. TCR + CD4 curve was fitted while keeping the same TCR affinity and off-rate as in the bimolecular interaction. p = 0.0552, 0.0162, 0.0624, 0.1179, 0.0523, and 0.0473 for comparing measured whole bonds to predicted sum at *t*_c_ = 0.5, 1, 2, 4, 8, and 16 s. **c,** Mean ± SEM bond lifetime vs force data of TPI-pulsed IFN-γ treated THP-1 cells interacting with beads presenting E8 TCR (n = 201), CD4 (n = 200), or E8 TCR and CD4 of similar density (n = 225). **d,** *P*_a_ vs *t*_c_ data (Mean ± SEM of 5 or more cells per point) and their model fits (curves) of pMHC-coated beads binding to Jurkat cells expressing TCR, CD4, or both at densities (μm^−2^) indicated by the legends. The dashed curve represents predicted sum of two concurrent and independent TCR–pMHC and CD4–MHC bimolecular interactions, which used the kinetic parameters of the two bimolecular interactions with molecular densities matched those used in the experiment measuring total interactions. Best-fit *P*_a_ ± SE was calculated from best-fit <*n*> ± SE in **e**. p = 0.0016, 0.0004, 0.0013, 0.00005, 0.0004, and 0.0066 for comparing measured *P*_a_ of whole interactions to predicted sum at *t*_c_ = 0.5, 1, 2, 4, 8, and 16 s. **e,** Normalized whole bond number <*n*>_W_/*m*_Receptor_*m*_pMHC_ vs *t*_c_ data (points) and their respective model fits (curves) for interactions in **d**. Predications were made after normalizing the densities in different curves. TCR + CD4 curve was fitted with (brown) or without (black) keeping the same TCR affinity and off-rate as in the bimolecular interaction. p = 0.0022, 0.0006, 0.0019, 0.0001, 0.0011, and 0.0140 for comparing measured whole bonds to predicted sum at *t*_c_ = 0.5, 1, 2, 4, 8, and 16 s. p = 0.0044, 0.0013, 0.0066, 0.0029, 0.3089, and 0.5179 for comparing measured whole bonds to its fitted curve (brown) at *t*_c_ = 0.5, 1, 2, 4, 8, and 16 s. **f,** Mean ± SEM bond lifetime vs force data of BFP beads bearing TPI:HLA-DR1 interacting with Jurkat cells expressing E8 TCR (n = 598), CD4 (n = 283), E8 TCR and CD4 of similar density (n = 279) or more TCR than CD4 (n = 288) expressed on Jurkat cells. **g,** Mean ± SEM bond lifetime vs force data of BFP beads bearing Hb:I-E^k^ interacting with 3.L2 CD8^+^CD4^−^ T cells (n = 298), CD8^−^CD4^+^ T cells (n = 711), or CD8^−^CD4^+^ T cells with CD4 blocking antibody clone GK1.5 (n = 240). **h,** Comparison among normalized bond numbers for pMHC interaction with TCR or mixed TCR and CD4 coated in four ways at densities (# of molecules/μm^2^) indicated by the number in legend: 1) pMHC on RBCs vs TCR on RBCs (green), 2) pMHC on RBCs vs mixed TCR and CD4 on RBCs (black), 3) pMHC on RBCs vs mixed TCR and CD4 on beads (orange), 4) pMHC on beads vs mixed TCR and CD4 on RBCs (blue). Data (points) are presented as mean ± SEM of two independent experiments for each curve. Their model fits (mean ± SE curves) are calculated using Eqs. 1 and 2 based on the best-bit parameters listed in Table 1.

When tested against beads coated with both TCR and CD4, the *P*_a_ (Fig. 3a, black) and the corresponding normalized whole bond number <*n*>_W_/*m*_TCR_×*m*_pMHC_ (Fig. 3b, black) were much higher than the predicted mediated by the sum of TCR–pMHC and CD4–MHC interactions calculated using their measured kinetic rates and molecular densities (Figs. 3a&b, dashed curves), demonstrating that purified TCR and CD4 cooperatively bind cellular pMHC. Interestingly, fitting the dual species data to Eqs. 1&2 while holding the TCR–pMHC and CD4–MHC kinetic rates the same values as in their bimolecular interactions yielded a good fit of the data with 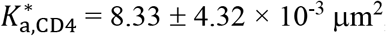 μm^2^, which is 0.16 times of that obtained using purified pMHC (Figs. 2a&b). Assuming a contact area of 1 μm^2^, the affinity of CD4 binding to TCR-engaged pMHC is ~1,823 fold of that for free pMHC. This fold increase is smaller than that obtained using purified pMHC and is likely due to the fact that in this case TCR and CD4 are immobilized on BFP beads instead of being freely diffusible on the RBC membrane.

To further confirm TCR-CD4 cooperativity for cellular pMHC, we measured bond lifetime vs force curves, finding that THP-1 expressed TPI:HLA-DR1 formed a slip-only bond, weak catch-slip bond, and strong catch-slip bond with beads coated with CD4 only, E8 TCR only, and both E8 TCR and CD4, respectively (Fig. 3c), consistent with the results obtained using purified pMHC (Fig. 2e).

Next, we analyzed purified pMHC binding to cellular TCR and CD4, and quantified the impacts of their transmembrane and cytoplasmic domains and of the CD3 signaling chains, which the purified TCRαβ and CD4 ectodomains lack and have been shown important in our previous studies of TCR-CD8 cooperativity^3,5^, on TCR-CD4 cooperativity. We performed experiments using Jurkat cells transfected with, and sorted to express comparable levels of E8 TCR, CD4, or both. We first analyzed the *P*_a_ vs *t*_c_ curves of these Jurkat cells binding to TPI:HLA-DR1 coated bead using the BFP setup (Fig. 3d). Comparing to purified TCRαβ coated on RBCs, cellular TCR-CD3 complex exhibits a ~1 log lower effective 2D affinity (*A*_c_*K*_a,TCR_ = 3.59 ± 0.37 × 10^−5^ μm^4^). In contrast, despite the smaller functional contact area, cellular CD4 has an effective 2D affinity (*A*_c_*K*_a,CD4_ = 6.57 ± 0.64 × 10^−6^ μm^4^) for purified pMHC that is ~1 log higher than that of purified CD4 for purified pMHC and similar to that of purified CD4 for cellular pMHC (Table 1).

Similar to the cell-free system, measurements with cells expressing both TCR and CD4 showed significantly higher adhesion frequency *P*_a_ (Fig. 3d, black) and normalized whole bond number <*n*>_W_/*m*_TCR_×*m*_pMHC_ (Fig. 3e, black) than the predicted the predicted sum mediated by the sum of TCR–pMHC and CD4–MHC interactions calculated using their measured kinetic rates and molecular densities (Fig. 3d&e, dashed curve), demonstrating that cellular TCR and CD4 cooperatively bind purified pMHC.

Comparing the predicted curve of Eqs. 1&2 with the <*n*>_W_/*m*_TCR_×*m*_pMHC_ vs *t*_c_ measurement showed excellent agreement between the model and data in both the logarithmic transformed variable (Fig. 3e) and the directly measured variable (Fig. 3d). Fitting the data returned values of 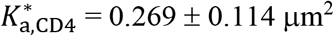 and *k*_−2,CD4_ = 0.131 ±0.117 s^−1^, respectively ~5 times of and ~1 log smaller than the corresponding values estimated from the cell-free system (Table 1). Interestingly, the values of *A*_c_*K*_a,TCR_ = 1.42 ± 0.463 × 10^−4^ μm^4^ and *k*_−1,TCR_ = 1.60 ± 1.06 s^−1^ estimated from fitting the <*n*>_W_/*m*_TCR_×*m*_pMHC_ data are, respectively, 4 and 4.7 times of the counterpart values estimated from fitting the respective <*n*>_TCR_/*m*_TCR_×*m*_pMHC_ and <*n*>_CD4_/*m*_TCR_×*m*_pMHC_ data. This is in contrast to both the results obtained by fitting the purified TCR-CD4 vs purified and cellular pMHC data where the corresponding numbers match (Table 1). Indeed, fitting the same whole bond data by Eqs. 1&2 using the *A*_c_*K*_a,TCR_ and *k*_−1,TCR_ values estimated from single bond species measurements as given rather than free fitting parameters resulted in a poor fit of significantly slower kinetics than the actual data (Fig. 3e brown curve), in sharp contrast to the excellent fits in Figs. 2b&3b where the <*n*>_W_/*m*_TCR_×*m*_pMHC_ curve and the <*n*>_TCR_/*m*_TCR_×*m*_pMHC_ curve have common initial slope at *t*_c_ = 0, as predicted by Eqs. 1&2. To compensate for the pre-determined lower *A*_c_*K*_a,TCR_ and slower *k*_−1,TCR_ values, the two-parameter fit (Fig. 3e brown curve) required much higher affinity 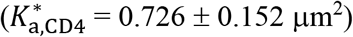 and off-rate (*k*_−2,CD4_ = 10.0 ± 16.9 s^−1^) for CD4 to respectively bind to and dissociate from TCR-prebound pMHC as compared to the four-parameter fit (Fig. 3e black curve). These results reveal similarities and distinctions between the cellular and purified TCR-CD4 systems and suggest that intracellular regulations, which are absent in the purified TCR-CD4 system but present in the cellular TCR-CD4 system, may speed up the first-stage kinetics to allow CD4 to cooperate in pMHC binding at an earlier time, a contention consistent with our previous results that intracellular regulations is important to the TCR-CD8 cooperative binding of class-I pMHC^3,5^.

Cellular TCR-CD4 cooperativity was also confirmed in bond lifetime analysis: TPI:HLA-DR1 formed a slip-only bond, weak catch-slip bond, and strong catch-slip bond with cells expressing CD4 only, E8 TCR only, and both E8 TCR and CD4, respectively, similar to their purified protein counterparts (Fig. 3f). In addition, Jurkat cells expressing a higher level of TCR than CD4 formed a more pronounced catch bond than those express similar levels of TCR and CD4 (Fig. 3f), suggesting a higher degree of cooperativity.

As an additional confirmation, we analyzed CD4^+^CD8^−^ and CD4^−^CD8^+^ single-positive T cells^32^ from 3.L2 TCR transgenic mice, which recognizes a hemoglobulin (Hb) epitope presented by mouse class-II pMHC I-E^k^ (Hb:I-E^k^)^8^. Our previous adhesion frequency experiment was unable to detect mouse CD4–Hb:I-E^k^ binding, indicating an *A*_c_*K*_a,CD4_ < 7.0 × 10^−8^ μm^4^ (the detection limit), no less than a log lower than the human CD4–HLA-DR1 effective 2D affinity, and observed a TCR–pMHC catch-slip bond between BFP probes bearing Hb:I-E^k^ and CD4^−^ CD8^+^ 3.L2 T cells by force-clamp BFP experiment^8^. Interestingly, when using CD4^+^CD8^−^ T cells we observed a much more pronounced catch-slip bonds, which was suppressed by anti-CD4 (GK1.5) blocking to the level of using CD4^−^CD8^+^ T cells (Fig. 3g). These data confirm that the prolonged lifetimes were CD4-dependent and further support the formation of more durable and force-resistant TCR–pMHC–CD4 trimolecular bonds by TCR-CD4 cooperation.

### TCR-CD4 cooperativity requires molecular mobility

The faster kinetics of the cellular TCR-CD4 cooperative binding to pMHC (Fig. 3e) may suggest enhanced CD4 recruitment to TCR–pMHC, a process that requires both molecular mobility and proximity. To test whether cooperativity depends on molecular diffusion, we used four different pairings of surfaces to present proteins: 1) pMHC on RBCs interacting with TCR only on RBCs; 2) pMHC on RBCs interacting with TCR and CD4 on RBCs; 3) pMHC on RBCs interacting with TCR and CD4 on beads; and 4) pMHC on beads interacting with TCR and CD4 on RBCs. The idea is to allow lateral diffusion of TCR and CD4 on the RBC membrane but prevent them from doing so on the bead surface. The data were presented as average number of whole bonds per unit densities of TCR and pMHC as in Fig. 2b. When interacting proteins on both sides were coated on RBCs (2^nd^ pairing), pMHC interaction with TCR-CD4 dual-receptor species (Fig. 3h, black) generated a much higher <*n*>_W_/(*m*_TCR_×*m*_pMHC_) curve than did with TCR single-receptor species (Fig. 3h, green), indicating cooperative binding as in Fig. 2b. In sharp contrast, when pMHC was coated on RBCs and TCR and CD4 were co-functionalized on beads (3^rd^ pairing), the binding curve was substantially downward shifted to the level indistinguishable to the curve generated by coating TCR-only on RBCs binding to RBCs bearing pMHC (1^st^ pairing, Fig. 3h, orange). This indicates that the cooperativity between TCR and CD4 vanished when they were immobilized at low densities on glass bead surface. When pMHC was coated on beads and the TCR/CD4 mixture was coated on RBCs (4^th^ pairing), the binding curve was upward shifted to a level higher than that resulted from coating proteins from both sides on RBCs, as predicted by the lower CD4 density (Fig. 3h, blue).

Using Eqs. 1&2 to globally fit the green and black data (Fig. 3h) returns *A*_c_*K*_a,TCR_ and *k*_−1,TCR_ values comparable to, and 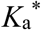 and *k*_−2,CD4_ values even larger than, those obtained by fitting Fig. 2b (Table 1). In sharp contrast, fitting the orange data (Fig. 3h) returns a vanishingly small 2D affinity of CD4 for TCR-prebound pMHC 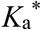 (1.0 × 10^−9^ μm^2^) with a 4-log larger standard error (4.2 × 10^−5^ μm^2^) despite reasonable values of *A*_c_*K*_a,TCR_, *k*_−1,TCR_, and *k*_−2,CD4_ (Table 1). Fitting the blue data (Fig. 3h) after constraining *k*_−1,TCR_, and *k*_−2,CD4_ to prevent over-fitting generated *A*_c_*K*_a,TCR_ and *K*_a^*^,CD4_ values similar to those obtained by globally fitting the green and black data (Table 1). Together, these results indicate that cooperative trimolecular interaction was modulated by the mobility of TCR and CD4 on the cell surface.

### Optimal TCR-CD4 cooperativity is achieved at 7 nm spacing

We hypothesized that the mobility effect might stem from the requirement for sparsely distributed TCR and CD4 to diffuse into close proximity for them to physically reach the same pMHC. To test this hypothesis, we designed a DNA nanostructure – a ~250-nm 10-helix bundle (10HB) DNA origami nanorod^33^, which can be used as a molecular ruler for precise control of the arrangement and relative locations of proteins^33^. 10HB nanorods functionalized with TCR and CD4 were captured on a magnetic bead to form a surrogate T cell (Fig. 4a) to test against a pMHC bearing RBC in the 2D kinetic assay (Fig. 4b). Many poly-A DNA strands protrude from one side of 10HB to bind poly-T strands on the bead surface, while TCR and CD4 are anchored by “capture” DNA strands on the other side of 10HB (Fig. 4c). The position of the capture DNA strands was changed to adjust the TCR-CD4 intermolecular spacing, which was designed to be 6, 13, 20, and 100 nm for this study (Fig. 4d). To bind the proteins to their complementary capture DNA strands on 10HB, CD4 was covalently linked with a DNA strand, verified by sodium dodecyl-sulfate polyacrylamide gel electrophoresis (SDS PAGE) (Fig. 4e). C-terminally biotinylated E8 TCR was bound to a streptavidin (SA), which was in turn bound to a biotinylated DNA strand on the 10HB (Fig. 4c). Successful assembly of 10HB nanorods was confirmed via native agarose gel electrophoresis (Fig. 4f). A slight decrease in mobility was observed after the addition of single-stranded extensions (poly-T strands and capture strands), compared to naked 10HB nanorods. The nanorods were extracted from the gel bands, purified, and used for attachment of TCR and CD4. Subsequent transmission electron microscopy imaging on negatively stained samples clearly showed the binding of TCR and CD4 and their 13-nm and 20-nm designed intermolecular spacings (Fig. 4g). We should note that the designed spacing are nominal, as the actual spacing can be affected by the length of DNA handles, the size of streptavidin, and the flexibility of the linkage.

**Figure 4.**
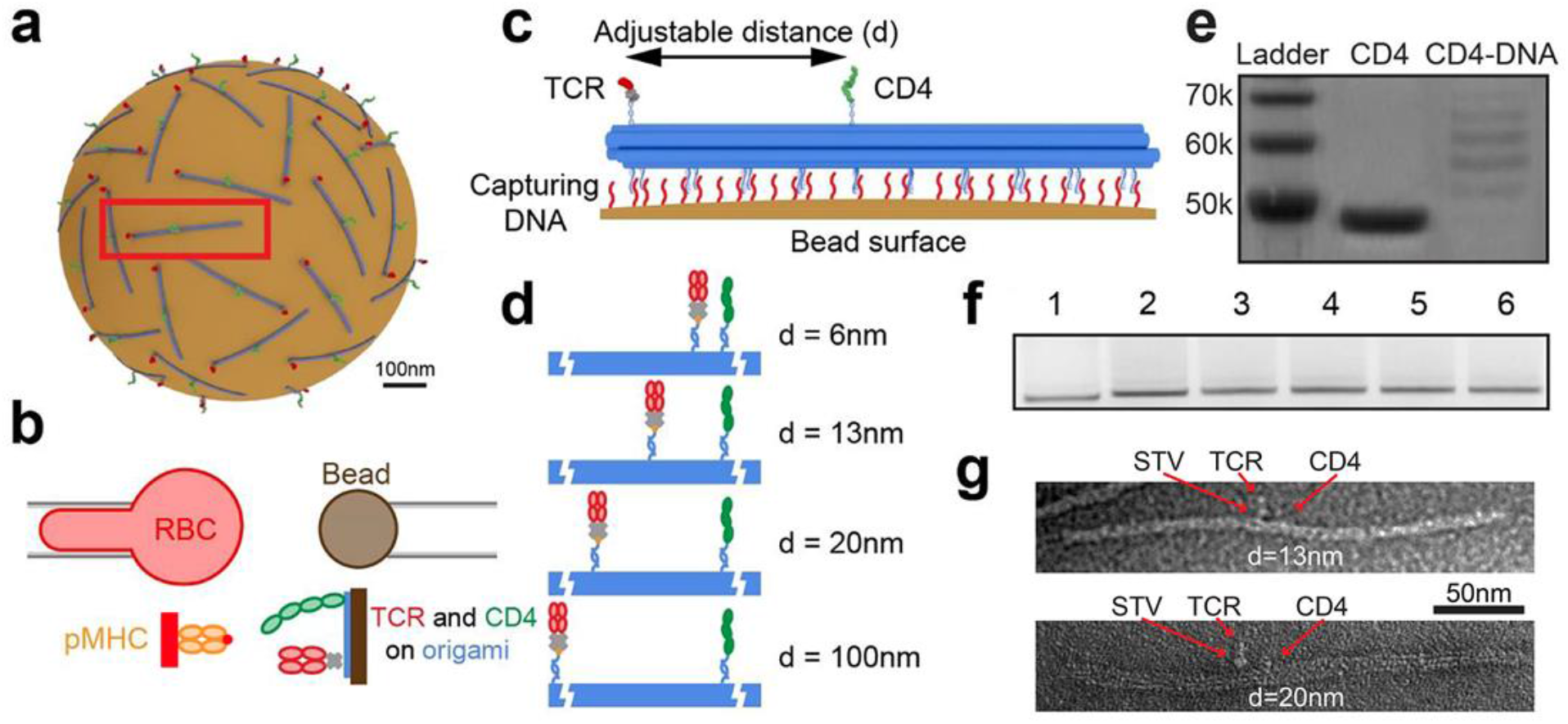
DNA origami-based platform for spacing control of TCR and CD4. **a,** Schematic of “surrogate” T cell – magnetic bead bearing 10 HB origami functionalized with precisely spaced TCR and CD4. **b,** Schematic of micropipette adhesion frequency assay combined with ligand presentation by origami. **c,** 10 HB origami controls separation distance of TCR and CD4 and attaches to magnetic bead via DNA strand hybridization. TCR binds to a streptavidin (SA), which also binds to a biotinylated DNA strand that hybridizes with a TCR-capturing DNA strand on 10HB origami. CD4 is covalently conjugated with a DNA stand that binds to a CD4-captruing strand. **d,** Four distances were employed, with either 6, 13, 20, or 100 nm TCR–CD4 spacing. **e,** SDS PAGE of CD4 conjugated to oligonucleotide for loading onto 10HB origami. **f,** Native agarose gel electrophoresis of 10 HB origami with 1) no poly-A extensions or protein handles, 2) poly-A extensions but no protein handles, 3-6) poly-A extensions with protein handles spaced at 6, 13, 20 and 100 nm respectively. **g,** TEM image of 10HB origami bearing CD4 and SA-coupled TCR with 13 nm (top) and 20 nm (bottom) spacing.

To examine how TCR-CD4 spacing impacted their synergistic binding to pMHC, we measured adhesion frequencies between RBCs bearing pMHC and origami beads coated with E8 TCR, CD4, or both with defined spacing. CD4-alone origami adhered negligibly as did protein/handle negative or origami negative controls, whereas the same density of TCR-alone origami yielded ~20% of *P*_a_ at steady-state (*t*_c_ = 4 s), consistent with its >3 logs higher affinity (Supplementary Fig. 4). Multiplexing TCR and CD4 on the origami 100 and 20 nm apart generated indistinguishable adhesions comparable to the TCR-alone group. Strikingly, further decreasing the spacing to 13 and 6 nm profoundly increased *P*_a_ (Supplementary Fig. 4) and its density-normalized log-transformation (Fig. 5a), indicating strong cooperativity as in Figs. 2b&3h.

**Figure 5.**
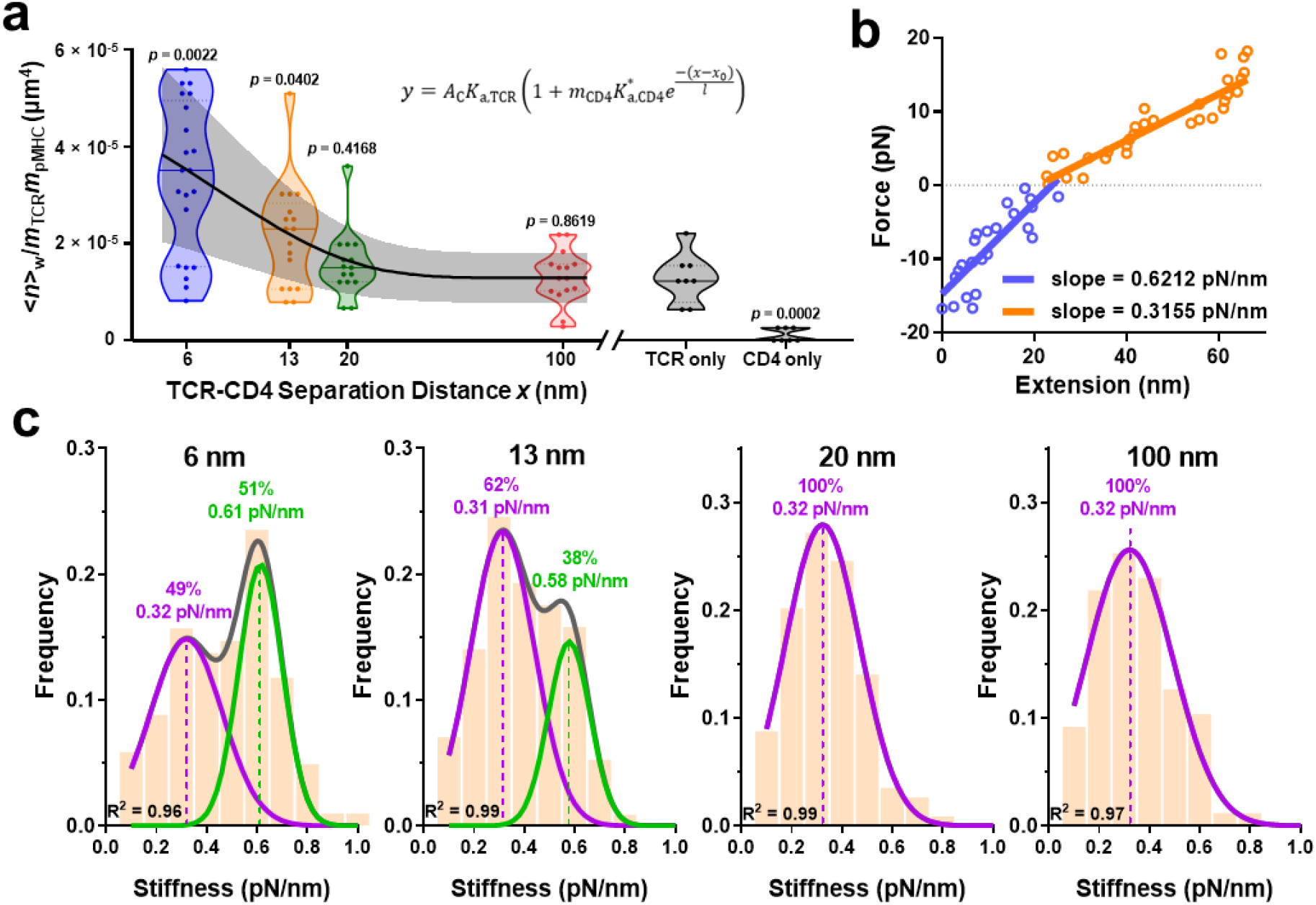
TCR-CD4 cooperativity is optimal at 7-nm spacing. **a,** Normalized whole number of bonds <*n*>_W_/*m*_TCR_*m*_pMHC_ for pMHC-bearing RBCs interacting with DNA origami beads presenting TCR-CD4 pairs at 6, 13, 20, or 100 nm spacing, or presenting TCR alone or CD4 along (points, n = 21, 17, 15, 15, 8, and 7 cell-bead pairs). In the violin plots that show data densities, the solid lines in the middle represent mean and the lower and upper dotted lines represent the first and third quartiles. Eq. 4 was fit (mean ± SE curves) to the <*n*>_W_/*m*_TCR_*m*_pMHC_ vs separation distance (*x*) data using the CD4 density (*m*_CD4_ = 31 μm^−2^) measured by flow cytometry. P-values indicate comparison with TCR only group using the Mann-Whitney test. **b,** Representative force vs molecular extension plot illustrating the determination of molecular stiffness using the Hook’s law. Blue: compression phase, where slope represents the spring constant of the cell. Orange: extension phase, where the slope represents the resultant spring constant of the cell the molecule connected in series. **c,** Histograms of stiffness of bonds between pMHC and TCR and CD4 spaced at 6, 13, 20, or 100 nm. Histograms were fitted by single gaussian distribution (20 and 100 nm) or double gaussian distribution (6 and 13 nm) to identify the peak value(s) and the associated fractions.

We extended Eq. 3 by assuming that increasing separation between TCR and CD4 would decrease the trimolecular bond formation exponentially, obtaining a phenomenological model that fits the data well:

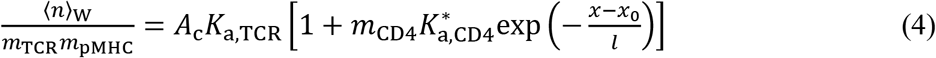

where *A*_c_*K*_a,TCR_ (1.29 ± 0.52 × 10^−5^ μm^4^) was obtained from the TCR-alone measurement. Its smaller than other values in Table 1 may be explained by presenting TCR on different surfaces (RBCs vs origami-coated beads)^34^. Fitting returned an apparent separation *x*_0_ (7.0 ± 1.7 nm) required for optimal cooperativity, decay length scale *l* (7.7 ± 2.1 nm), and CD4 affinity for TCR pre-engaged pMHC 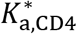 (0.050 ± 0.015 μm^2^). The *x*_0_ value agrees well with the 7-nm gap between the membrane anchors of TCR and CD4 found in their ternary structure complexed with pMHC^12^, and values <7 nm lead to worse fits. The *l* value describes how fast the ability of CD4 to bind TCR-engaged pMHC would decay exponentially as *x* increases beyond *x*_0_, underscoring the requirement for spatial proximity. The 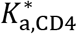 value is statistically indistinguishable from the value estimated from Fig. 2b using Eqs. 1&2 (Table 1).

To obtain independent support for the existence of trimolecular bonds, we analyzed single-bond stiffness to distinguish bimolecular vs trimolecular interactions^5^ (Fig. 5b&c). When TCR and CD4 were spaced 20 or 100 nm apart, the histograms of molecular stiffness exhibited a single peak well fitted by a Gaussian distribution, yielding a mean stiffness of 0.32 pN/nm for both samples (Fig. 5c). Spacing the molecules 13 nm apart gave rise to a double-peak histogram (Fig. 5c). The first peak has nearly the same mean stiffness of 0.31 pN/nm but the second peak represents a subpopulation (38% of total) of nearly doubling the stiffness (0.58 pN/nm), indicating the presence of trimolecular bonds. The 6-nm spacing further populates the second peak (51%) but maintains the mean stiffness values for the two peaks (Fig. 5c). Interestingly, these fractions are consistent with the corresponding fractions with low and high average numbers of whole bonds (<*n*>_W_) normalized by the TCR and pMHC densities (*m*_TCR_ and *m*_pMHC_) measured at 13 and 6 nm using the 2D assay (Fig. 5a). Overall, the data indicate that a threshold distance is required for TCR and CD4 to form trimolecular bonds with pMHC.

## Discussion

A previous study using FRAP to analyze the mobility of TCR-CD3 or CD4 at the interface with pMHC observed reduced mobility of CD4 in the presence of TCR–pMHC interaction and reduced TCR-CD3 mobility in the presence of CD4–MHC interaction^22^, suggesting the binding of cellular TCR and CD4 to the same pMHC. Using two ultrasensitive mechanical assays to measure adhesion frequencies and single-bond lifetimes of three related 2D interaction systems – purified TCRαβ and CD4 ectodomains vs purified pMHC ectodomain, purified TCRαβ and CD4 ectodomains vs cellular pMHC, and cellular TCR-CD3 and CD4 vs purified pMHC ectodomain – and presenting the soluble proteins on the surface of RBCs, glass beads, and DNA origami, we showed in detail how and why this occurs despite the extremely low CD4–MHC affinity. By demonstrating that CD4 binds TCR-prebound pMHC with 3-6 logs (depending on the system) higher affinity than it binds free pMHC, our results explain how CD4 could efficiently augment TCR antigen recognition via stabilizing TCR–pMHC bonds and recruiting Lck to TCR-CD3. This also answered a perplexing question of why anti-CD4 blocking antibody would suppress Lck phosphorylation of CD3^35^, if the CD4 could not even bind pMHC. While CD4 alone cannot initiate T cell interaction with APC, it is able to provide adequate “help” to TCR in recognition of cognate pMHC via enormous synergy and cooperation, a mechanism that greatly enhances TCR sensitivity.

Our work highlights the regulatory effects of force on dynamic cooperative molecular bonds. The cooperativity of mouse CD4 and TCR binding to Hb:I-E^k^ could only be detected by the BFP force-clamp assay but not by the force-free adhesion frequency assay^8^, suggesting that force on pMHC may enhance its on-rate for CD4. Our model also suggests that force-elicited catch bond enhances the sustainability of the pMHC–CD4 arm of the trimolecular bond by prolonging its lifetime.

An intriguing question raised by our findings is the molecular basis of the enhanced affinity of CD4 for TCR-prebound pMHC. The present work has ruled out the possibility that the TCR-CD4 cooperativity requires TCR–pMHC binding to induce inside-out signaling of CD4 to upregulate its affinity for pMHC, since soluble TCRαβ and CD4 proteins coated on RBC, glass bead, and DNA-origami surfaces are capable of cooperative binding to purified and cellular pMHC. Another possibility is that TCR binding to pMHC induces conformational changes in the CD4 binding site of MHC that increase affinity for CD4. However, crystallographic studies of numerous TCR and pMHC molecules in free and bound form provide no evidence of TCR-induced structural changes at the site on the MHC class II β2 domain where CD4 binds. This would argue against an allosteric mechanism to explain increased CD4 affinity for TCR-prebound MHC. However, it is also possible that the relevant changes may be in protein dynamics, a parameter that cannot be addressed via the static snapshots provided by X-ray crystallography. Indeed, several NMR studies have found that pMHC binding to TCR induces long-range allosteric changes in TCR dynamics that could be relayed to CD3, even in the absence of obvious structural changes in the TCR^36–38^. However, the reverse situation (i.e. long-range allosteric changes in pMHC dynamics upon TCR binding) has not been examined by NMR, but probably should be given our results here.

The proposed structural mechanisms are supported by our modeling analysis, which was used to fit the bond lifetime data measured by the BFP experiments (Figs. 2d-f, Supplementary Fig. 3). To show this, we first compared the abilities of three models to simultaneously fit the three specific interaction data measured by thermal fluctuation (Fig. 2d). The first model is depicted in Supplementary Fig. 3a except that the activation to and dissociation from the strong state (TCR–pMHC)* is neglected because force was zero. Also, the weak (TCR–pMHC–CD4) and strong (TCR–pMHC–CD4)* trimolecular bonds are assumed to be in equilibrium and the two states dissociate concurrently to free molecules TCR + pMHC + CD4 in a single step (hence called concurrent model). The second model is a single-pathway, two-step, sequential dissociation model (sequential model I). It differs from the concurrent model in that the TCR–pMHC–CD4 trimolecular bond has only a single state that dissociates sequentially from right to left in Eq. 9 (Methods), first the pMHC–CD4 arm reversibly to generate TCR–pMHC complex + CD4, and then the TCR–pMHC bimolecular bond to free molecules TCR + pMHC + CD4. The third model is also a single-pathway, two-step, sequential dissociation model (sequential model II). It is similar to the sequential model I except that the order of sequential dissociation of the TCR–pMHC–CD4 trimolecular bond is reversed, which occurs first at the TCR–pMHC arm reversibly to generate pMHC–CD4 complex + TCR, and then at the pMHC–CD4 bimolecular bond to generate free molecules TCR + pMHC + CD4. Using the Akaike Information Criterion (AIC) to choose between two non-nested models, we found that the concurrent model is more likely to be correct than sequential model I by >58,000 times and even much more times than sequential model II. This conclusion still holds even when we analyzed data measured by force-clamp assay (Supplementary Fig. 3b) to evaluate their force-dependent kinetic parameters (Supplementary Fig. 3c,d). The concurrent model suggests a strong functional coupling between the spatially separate TCR binding site and the CD4 binding site on the pMHC, supporting our allostery hypothesis. Future studies are required to test this hypothesis. Moreover, to confirm that association and dissociation happen preferentially via pathway I instead of pathway II, we considered a more general two-pathway, two-step model that combines both sequential models I and II, and globally fit the adhesion frequency data from Fig. 2a-b. We compared it to the sequential model I by AIC, and found little difference statistically, suggesting that both association and dissociation along the first pathway (i.e., sequential model I) dominates over the second pathway (i.e., sequential model II), which only happens with very low probability (~0.99 and ~0.01 for pathway I and II, respectively, of the two-pathway, two-step model).

This work builds upon our previous findings of TCR-CD8 cooperativity for class-I pMHC binding^3–5,7,20^ and adds to our previously proposed mechanistic model^5,39^. In live-cell systems, cooperativity relies on the recruitment of coreceptor to phosphorylated CD3 via a Lck “bridge”, where both the adaptor function and kinase function of Lck are required^3,5,20,39,40^ as seen with co-immunoprecipitation studies that identify similar molecular assembly connecting CD4 or CD8 to triggered TCR-CD3^15,41–44^. Here we showed that using purified TCR and CD4 lacking cytoplasmic domains and CD3, the synergistic cooperation could still occur if both molecules are allowed to diffuse on the RBCs or packed closed enough on the bead surface, suggesting that modulation of ectodomain cooperation could be decoupled from intracellular regulation. This is also consistent with the observation that CD4 lacking the ability of binding Lck is still required for TCR-mediated IL-2 production^22^. On the other hand, our data of cellular TCR and CD4 cooperating with a faster kinetics than their purified ectodomain counterparts regardless of being tested against purified pMHC or cellular pMHC suggest that there may be cellular regulation(s) to expedite TCR-CD4 cooperativity. Modulation of ectodomain by physical factors such as mobility and colocalization are intuitive, but require 2D analyses to decipher. Indeed, by limiting mobility using different surface and controlling TCR-CD4 proximity using DNA origami, we found a 7-nm apparent spacing for TCR-CD4 cooperativity, such that increasing the spacing resulted in an exponential decay in the capacity to form TCR–pMHC–CD4 trimolecular bonds. Such physical regulatory mechanisms revealed by the in vitro system further advance our understanding of how interactions between membrane receptors and ligands critically depend on their cell surface organization. Methodologically, our DNA origami-based 2D kinetic assays hold great potential to dissect the spatial requirement of the cooperation of single- or multi-receptor–ligand species by ectodomain binding and/or intracellular crosstalk.

## Methods

### Proteins

Biotinylated CD4 and biotinylated αβ E8 TCR were prepared as previously described^15,19^. Briefly, a 17-amino acid tag (TPI, GGGLNDIFEAQKIEWHE) was added to the C-terminus of CD4 ectodomain and to the C-terminus of the E8 TCRα ectodomain. CD4 was expressed in baculovirus-infected Sf9 insect cells^15^. E8 TCR was produced by *in vitro* folding from inclusion bodies expressed in *E. coli*^19^. The purified proteins were biotinylated using biotin protein ligase (Avidity); excess biotin and ligase were removed with a Superdex 200 column (GE Healthcare). The control MHC-I (HLA-A2) presenting the melanoma antigen peptide NY-ESO-1157-165 (9V, SLLMWITQV) was produced by the NIH Tetramer Facility at Emory University.

Biotinylated TPI: HLA-DR1 was made by the NIH Tetramer Core facility (Atlanta, GA) with peptide (GELIGTLNAAKVPAD) purchased from GenScript. Divalent streptavidin was generously gifted by Dr. Baoyu Liu (University of Utah).

### Cells

Human RBCs were isolated from blood of healthy donors according to a protocol approved by the Institutional Review Board of Georgia Institute of Technology. For adhesion frequency assay, RBCs were purified by Histopaque-1077, washed with ice cold PBS, and resuspended in EAS-45 buffer (2 mM Adenine, 110 mM D-glucose, 55 mM D-Mannitol, 50 mM Sodium Chloride, 20 mM Sodium Phosphate, and 10 mM L-glutamine). Equal aliquots of RBCs were then mixed with various concentrations of EZ-Link Sulfo-NHS-LC-Biotin (Thermo Scientific) at a pH of 7.2 for 30 min at room temperature, yielding different densities of biotin sites on RBC surfaces. Biotinylated RBCs were washed with EAS-45 buffer and stored at 4°C. For BFP experiments, freshly isolated human RBCs were biotinylated with biotin-PEG3500-NHS (Jenkem Technology) and then incubated with nystatin in N2 buffer (265.2 mM KCl, 38.8 mM NaCl, 0.94 mM KH2PO4, 4.74 mM Na2HPO4, and 27 mM sucrose; pH 7.2 at 588 mOsm) for 30 min on ice. Nystatin-treated biotinylated RBCs were washed twice with N2 buffer and stored at 4°C for BFP experiments.

HEK 293T cells were purchased from ATCC (Manassas, VA) and cultured in DMEM supplemented with 10% FBS, 6 mM L-glutamine, 0.1 mM MEM non-essential amino acids, and 1 mM sodium pyruvate. TCR β-chain deficient Jurkat J.RT3 cells were purchased from ATCC (Manassas, VA) and cultured in RPMI 1640 supplemented with 10% FBS, 100 U/mL penicillin, 100 μg/mL streptomycin, 2 mM L-glutamine, and 20 mM HEPES. J.RT3 were transduced by lentivirus to express the E8 TCR. Briefly, E8 TCR α and β chains joined by a P2A element were subcloned into pLenti6.3 vector with a T2A-rat CD2 reporter. Lentivirus encoding E8 TCR were produced by co-transfection of HEK 293T cells with E8TCR-pLenti6.3, pMD2.G (Addgene #12259), and psPAX2 (Addgene #12260) using lipofectamine 3000 (ThermoFisher Scientific). J.RT3 cells were transduced by incubating overnight with supernatant containing lentivirus and FACS sorted using Aria cell sorter (BD Biosciences) based on the surface expression E8 TCR. E8TCR J.RT3 cells were then transfected using Nucleofection (Lonza, Morristown, NJ) to express full-length human CD4 and FACS sorted using Aria cell sorter (BD Biosciences) for stable CD4 expression. THP-1 cells were cultured in RPMI 1640 supplemented with 10% FBS, 100 U/mL penicillin, 100 μg/mL streptomycin, 2 mM L-glutamine, and 20 mM HEPES. One day prior to experiment, the cells were treated with 20 – 100 U/ml IFN-γ (R&D Systems) and 1 μM TPI peptide (Genscript).

### Site density measurements

Site densities of TCR, CD4, and pMHC on RBCs, Jurkat cells, THP-1 cells, glass beads, or origami beads were measured by flow cytometry using PE-conjugated antibodies from BD Biosciences (San Jose, CA): anti-human TCR-β1 (clone JOVI.1, 1:20), anti-human CD4 (clone OKT4, 1:20), and anti-human HLA-DR (clone L243, 1:5). Isotype control antibodies were PE mouse IgG2a κ (clone G155-178, 1:20) for TCR and HLA-DR, and PE mouse IgG2b, κ (Clone 27-35, 1:20) for CD4. RBCs or beads were incubated with corresponding antibodies in 1X PBS + 2% BSA for 30 min at 25 °C and washed three times before being analyzed using the BD Accuri Flow Cytometer. Site density was calculated based on antibody-conjugated PE fluorescence relative to the QuantiBRITE PE standard beads (BD Biosciences). For micropipette experiments, RBCs bearing protein of interest (TCR, CD4, or MHC) were prepared for flow cytometry using three separate samples: one saturating the RBC surface, one of the experimental conditions, and one isotype control. Brightness across all antibodies was normalized against the CD4 staining which was brighter than the TCR staining.

### Micropipette Adhesion Frequency Assay

The theoretical framework and detailed experimental procedures of this assay have been described previously^23,24,45^. This assay leverages the ultra-soft RBC membrane to detect interactions with single-bond sensitivity. Briefly, RBCs with controlled biotin sites were first incubated with 200 μg/mL divalent streptavidin and washed, followed by subsequent coating with biotinylated pMHC, TCR, CD4, or a mixture of TCR and CD4. After washing, RBCs coated with receptor-ligand pairs were injected into an imaging chamber containing non-CO2 dependent L15 media supplemented with HEPES and 2% BSA. During experiment, two RBCs were aspirated by micropipette and repeatedly brought into contact of defined duration by a program-controlled piezo. Adhesion between two RBCs were detected by the deflection of RBC membrane during the separation after each contact with ‘1’ indicating an adhesive event and ‘0’ indicating no adhesion. Each pair of RBCs were tested for 50 cycles of the same contact duration to yield the adhesion frequency (*P*_a_), which is as the average score from the 50 cycles. The same process was repeated for multiple pairs of RBC of each contact duration (*t*_c_), yielding a *P*_a_ vs *t*_c_ curve that is fitted to a published model^23,24^ (see below).

Non-specific controls were conducted for all sets of species coatings. To do so, RBCs coated with divalent streptavidin anchored irrelevant protein (OT1 TCR) were brought into contact with RBCs coated with proteins of interest. An adhesion frequency curve would be obtained for nonspecific adhesion. Then, for each contact time, the distribution of nonspecific adhesion frequency would be removed from the measured distribution of adhesion frequency^27–29^

In some experiments, one or both sides of the RBCs were replaced by glass beads coated with pMHC (see below) or THP-1 cells expressing pMHC, or glass beads or origami beads coated with TCR, CD4, or both (see below), or expressed on Jurkat cells expressing TCR, CD4, or both.

### Biomembrane Force Probe Assay

The principle and experimental procedures for BFP assays have been described previously^11^. To coat proteins on beads, borosilicate glass beads were mercapto-propyl silanated and covalently functionalized to monovalent streptavidin-maleimide in phosphate buffer saline (pH 6.8) by overnight incubation at 25°C. Streptavidinylated beads were then incubated with sub-saturating concentrations of biotinylated pMHC, TCR, CD4, or 1:1 mixture of TCR and CD4 for 2 hours in HBSS + 2% BSA at 4°C. During the experiment, a protein-coated bead (tracking bead) was attached to the apex of a micropipette-aspirated RBC that serves as a spring with pre-adjusted spring constant of 0.1-0.3 pN/nm. For force clamp assay, the tracking bead was repeatedly contacted by a piezo-driven target bead coated with (or target cell expressing) the corresponding binding partner(s). The displacement of tracking bead was monitored with a high-speed camera at 1000 fps and was translated into force by applying the pre-defined spring constant. During separation, bond formation between tracking bead and target bead/cell pulled the tracking bead away from its baseline, manifesting positive force loading on the molecular bond. Bond lifetime was defined as the duration from the start of clamp at the preset force level to bond rupture. Several hundred bond lifetime events were collected and pooled for various clamp forces using multiple bead-bead or bead-cell pairs. The stiffnesses of molecular bonds were modeled as Hookean spring constants which is determined by the difference between the displacements of the bead tracking system (*F*) and the displacements of the piezoelectric actuator-capacitance sensor feedback system (Δ*x*). Two straight lines were fitted to the piecewise data: One was fitted to the compressive force regime where the bead was slowly ramping away from cell impingement (force < 0), where the slop represents the spring constant of the cell membrane (compression). The other was fitted to the tensile force regime where the bead would ramp away from the cell beyond the point of initial contact (force > 0), where the slop represents the resultant spring constant of cell membrane (extension) and molecular bond connected in series. By assuming the spring constant of cell membrane is identical during compression as during extension, the spring constant of molecular bond was extracted by applying Hooke’s law for springs in series. An ensemble of molecular spring constant was collected as histograms using similar force bins and fitted to single or double Gaussian distributions.

For thermal fluctuation assays^24,30^, a target bead was driven by piezo to the proximity (~10 nm) of the probe bead, allowing bond formation between the two during intermittent contacts caused by thermal fluctuation, which would decrease the fluctuation of probe bead position. By analyzing the standard deviation (SD) of the tracking bead position, bond formation and rupture were identified by the decrease of SD below a cutoff of 3 nm and the subsequent resumption SD to baseline level. The cutoff baseline was determined as the lowest SD value of the probe bead before contact (*i.e.*, SD of bead without bonds).

For adhesion frequency assays using BFP beads, the same process of RBC-based adhesion frequency assay was performed except that a target cell was repeatedly brought into contact with a pMHC-coated bead, held for a defined duration, and then separated. Adhesion was defined as the deflection of bead away from its equilibrium position.

### Kinetic models

#### Single-step model for bimolecular interaction

The bimolecular interactions of either TCR or CD4 with pMHC are modeled as a single-step reversible reaction:

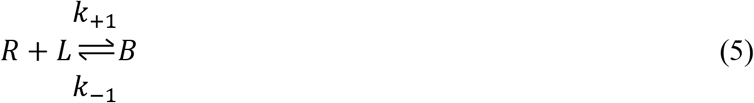

where *R, L,* and *B* denote, respectively, receptor, ligand, and bond; *k*_+1_ and *k*_−1_ denote, respectively, on- and off-rates. Their ratio is the binding affinity, *K*_a_ = *k*_+1_/*k*_−1_. Our published probabilistic kinetic model for the above reaction states that the probability of adhesion (*P*_a_) depends on the contact time (*t*_c_) and area (*A*_c_) as well as the densities of receptors (*m*_r_) and ligands (*m*_l_)^23,24^.

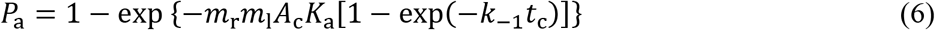

On the right-hand side, the term inside the braces after the minus sign is the average number of bonds per contact <*n*>.

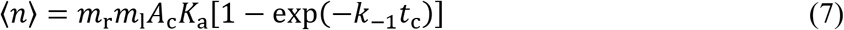

After normalizing <*n*> by *m*_r_*m*_l_, the right-hand should be independent of the molecular densities and approaches *A*_c_*K*_a_, the effective 2D affinity, as *t*_c_ becomes large.

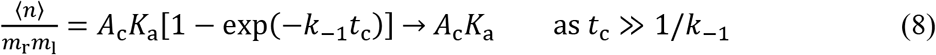

It follows from Eq. 8 that <*n*> is proportional to *m*_r_*m*_l_ if measured at large contact time.

We use TCR or CD4 in the subscript to identify the interaction with which the kinetic parameters are associated, e.g., *k*_+1,TCR_, *k*_−1,TCR_, and *K*_a,TCR_ are kinetic rates and binding affinity for the TCR–pMHC interaction, which are different from *k*_+1,CD4_, *k*_−1,CD4_, and *K*_a,CD4_, which are corresponding parameters for the CD4–MHC interaction. Also, *m*_TCR_, *m*_CD4_, and *m*_pMHC_ are used to designate the densities of TCR, CD4, and pMHC.

#### Two-step model for trimolecular interaction

We propose a two-step model for the formation of trimolecular bonds. Since the affinity and on-rate for the CD4–pMHC interaction are so much smaller than those of the TCR–pMHC interaction, it seems reasonable to assume that TCR interacts with pMHC at the same kinetic rates as bimolecular interaction in the first step as if CD4 does not interact with free pMHC. However, CD4 is able to interact with TCR-stabilized pMHC with a much higher affinity, thereby forming trimolecular bonds in the second step.

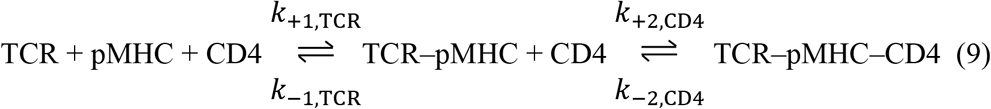

Let *p*_m,n_ be the probability of having *m* TCR–pMHC bonds and *n* TCR–pMHC–CD4 bonds, which are governed by the following master equations:

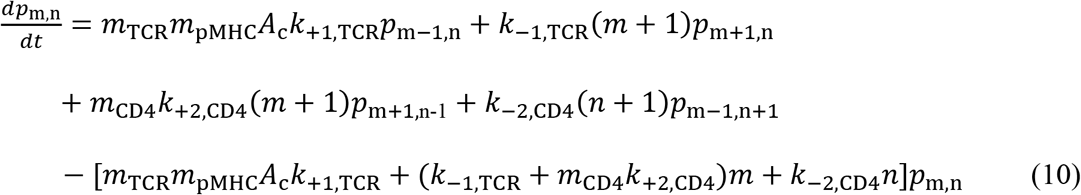

We solved these master equations using a probability generating function,

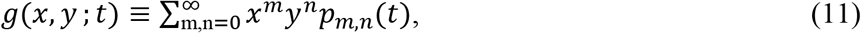

which converts Eq. 10 to a single first-order linear partial differential equation,

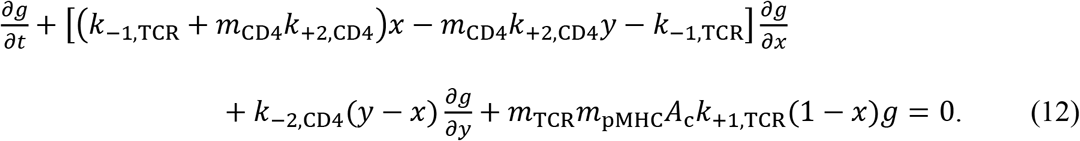

We found a general solution to Eq. 12 using the method of characteristics, which is

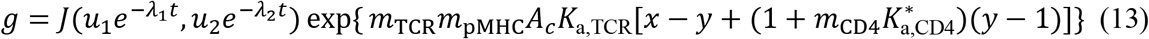

where *J* is an arbitrary function of its two arguments. *u*_1_ and *u*_2_ are functions of *x* and *y*:

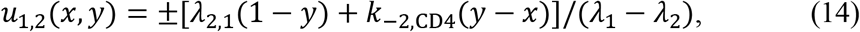

*λ*_1_ and *λ*_2_ are given by Eq. 2. They can be viewed as the fast (*λ*_1_) and slow (*λ*_2_) rates that control the two p44hases of the two-step interaction. To determine the function of integration *J* requires initial conditions. If there is no TCR–pMHC or TCR–pMHC–CD4 bonds at *t* = 0, the initial conditions on *p*_m,n_ are:

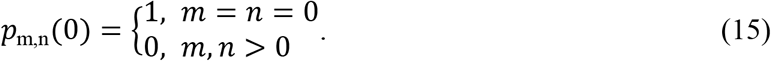

The corresponding initial condition of *g*, obtained by substituting Eq. 15 into Eq. 11, is

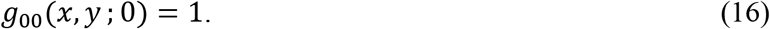

Solving *x* and *y* in terms of *u*_1,2_ from Eq. 14, substituting them into Eq. 16, and comparing the resulting equation to Eq. 13 evaluated at *t* = 0 allows us to solve for a particular solution for *J*:

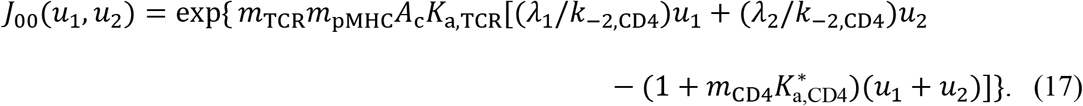

Substituting Eq. 17 into Eq. 13, we have

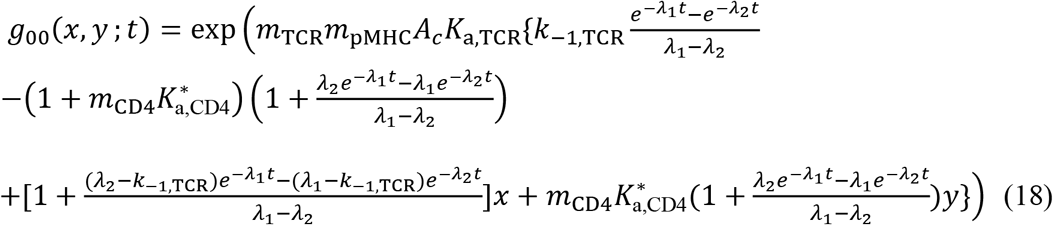

Expanding the right-hand side of Eq. 18 into Taylor series of *x* and *y*, and comparing the coefficients of the *x*^m^*y*^n^ terms to those of the right-hand side of Eq. 11, we found

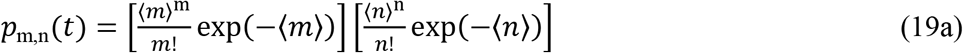

where

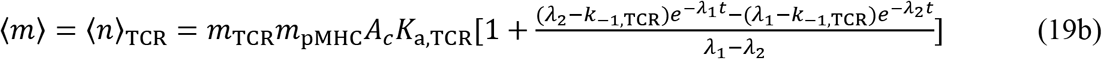

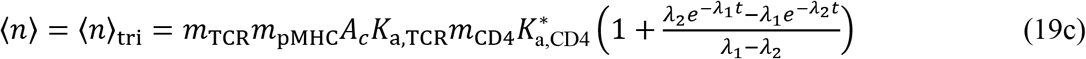

are the average numbers of TCR–pMHC bonds and TCR–pMHC–CD4 bonds. Thus, the solution is of the form of the product of two Poisson distributions. Except for the expressions for <*m*> and <*n*>, this is the same as the bond distribution of the dual species concurrent independent binding model described previously^27–29^.

The adhesion probability is the probability of having at least one bond of either type, or one minus the probability of having no bond, which can be found from Eq. 19:

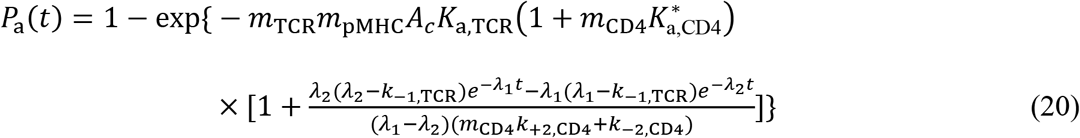

Taking a log transformation, we obtain the average number of whole bonds 〈*n*〉_W_ = – ln(1 – *P*_a_). Normalizing 〈*n*〉_W_ by the densities of TCR and pMHC, we obtain Eq. 1. Setting *k*_+2,CD4_ = *k*_−2,CD4_ = 0 reduces Eq. 2 to *λ*_1_ = *k*_−1_ and *λ*_2_ = 0, which also reduces Eqs. 20 and 1 to Eqs. 6 and 8, respectively, as expected.

### DNA Origami preparation

Origami were created at 20 nM using p8064 scaffold, 5× staple excess, and 25 × biotin strand for TCR capture. Following agarose gel electrophoresis in 1% agarose at 60 V for 90 min on ice bath, origami were purified with Freeze ‘n’ Squeeze spin columns. The four samples with different protein separation distances were indistinguishable based on gel mobility. However, a control sample with only handles for surrogate cell attachment ran with slightly increased mobility, while a complete blank 10 HB sample with no handle extensions whatsoever ran markedly faster through the gel. To confirm proper assembly of protein onto the DNA origami, purified samples were incubated with excesses of CD4–DNA and wild type streptavidin at 4 ^o^C overnight. After another agarose gel purification, a 10-fold molar excess of biotinylated TCR was added and allowed to incubate at 4 ^o^C for 3 h. Excess protein was removed using 100 kD MWCO spin filters, then the samples were deposited on 400 mesh carbon formvar grids and stained with 1% uranyl formate solution for 30 s. During TEM imaging, the identity of blinded samples could be reliably divined by observing relative CD4, SA–TCR location.

An amine-modified DNA strand was conjugated to lysine residues in CD4. CD4 was reacted with 50-fold excess succinimidyl-6-hydrazino-nicotinamide (S-HyNic) (Solulink) for 4 h, and terminal amine DNA was reacted with 20-fold excess succinimidyl-4-formylbenzene (S-4FB) (Solulink) for 4 h. Amicon Ultra spin filters (Millipore) were used to remove excess linker and buffer exchange into citrate buffer (50nM sodium citrate, 150 mM NaCl, pH 6). Functionalized DNA was combined with CD4 at a 3:1 ratio, and reacted overnight. CD4-DNA conjugates were purified on a Superdex 200 Increase 10/300 GL column using an AKTA Pure FPLC (GE Healthcare). Conjugation was verified with SDS/PAGE and Coomassie blue staining.

In order to examine the effect of geometric docking on the simultaneous interaction of two molecules with a third, origami structures differentially spacing TCR and CD4 were coated on beads to represent a simplified T cell surface. Beads were respectively coated with several different origami structures: those with handles for TCR and CD4 which were differentially spaced apart at 6, 13, 20, and 100 nm; those lacking protein handles; and origami lacking connecting strands to the beads. The purified samples were loaded onto surrogate T cells – 3 μm diameter magnetic dt Dynabeads. For each sample, 15 μL of beads were washed 3× with PBS then added to 1 μg (~30 μL) of purified DNA origami. A large excess of CD4–DNA and streptavidin were added, and the samples were left to incubate on a rotating shaker at 4 ^o^C overnight. The magnetically pelleted beads were washed with PBS supplemented with calcium and magnesium, and 2% BSA five times. Biotinylated (at the C-terminus of the α-chain) E8 TCR was added to the beads and incubated for 30 min. To this solution, blocking DNA strand (25 dA) was added and the reaction incubated at 4 °C for an additional 1 h.

### Fitting models to data

Linear fittings of data in Fig. 2d, Fig. 5b, Supplementary Fig. 1, and Supplementary Fig. 2c were implemented in Prism using linear regression to estimate the slope and *y*-intercept of the trend. Non-linear fittings of bimolecular kinetic equations to data in Figs. 1c,d were implemented in Prism using Non-linear regression to estimate the corresponding kinetic parameters. Gaussian fitting of data in Fig. 5c was implemented in Prism using non-linear regression to estimate the Gaussian distribution and frequency of each distribution. For all fitted parameters, fitting error was presented as Standard Error (SE) assuming symmetrical confidence intervals.

The model predicted curves in Figs. 2a-c and Supplementary Fig. 2a were obtained by globally fitting all data to Eqs. 1&2, which were also fit to data in Fig. 3a,b,d,e,h. For the Fig. 3h case, the green and black data were globally fit, the orange data was fit on its own, and the blue data was fit after constraining the off-rates obtained from the orange data fit to avoid overfitting. The separation distance-dependent adhesion frequency of the DNA origami assay (Fig. 5a) was fit to Eq. 4. For Figs. 2a-c, 3a,b,d,e,h, a two-step optimization procedure was employed to minimize the mean squared error (MSE) of the fits with respect to the data, which explored a large parameter space by Differential Evolution to obtain an approximate solution, followed by L-BFGS to hone in the minimum. The parameters Standard Errors (SE) were calculated from the covariance matrix obtained by inverting the Hessian matrix of the loss function at the minimum found. The SEs were then propagated through the model to display error bands of the fits as mean ± SE. The data in Fig. 5a were also fitted by the aforementioned two-step optimization procedure but the parameters SE were obtained via bootstrapping, with 1000 bootstrap samples.

The force-dependent lifetimes of pMHC bonds with TCR alone or co-presented TCR and CD4 behaved as catch-slip bonds, while pMHC bonds with CD4 alone behaved as a slip bond. Although at zero force the two catch-slip bonds had mono-exponentially decaying survival probabilities (Fig. 2d), a second population appeared at forces >0, displaying biexponentially decaying survival probabilities, suggesting the presence of two bound species, even for the bimolecular TCR–pMHC. Therefore, while the CD4–pMHC bonds were modelled to dissociate along a single pathway, both TCR–pMHC and TCR–pMHC–CD4 bonds were modelled to dissociate along two pathways – a fast dissociating pathway from a weak state or a slow dissociating pathway from a strong state (indicated by *) allowed to exchange from each other as depicted in Supplementary Fig. 3a (left and right panels) similarly to previous catch bond models^46,47^ The force dependence of all kinetic rates follows Bell’s equation^48^:

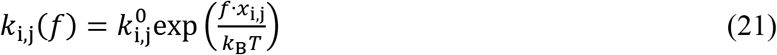

where *f* is force, the superscript 0 denotes the zero-force kinetic rate constant, the subscript i denotes rates of dissociation along fast (-f) or slow (-s) pathways and of activation (a) or deactivation (-a) transition between the weak and strong states, the subscript j denotes different bonds (TCR–pMHC or TCR–pMHC–CD4), *x*_i,j_ denotes distance to the transition state, or width of the energy well that traps the system in the bound state, *k*_B_ is Boltzmann constant, and *T* is absolute temperature. The weak state dissociates more rapidly than the strong state, so *k*_−f,i_ > *k*_−s,i_. The two-states, two-path catch-slip model can be expressed in terms of ordinary differential equations that govern the probability of having bonds *P* at a given time *t*:

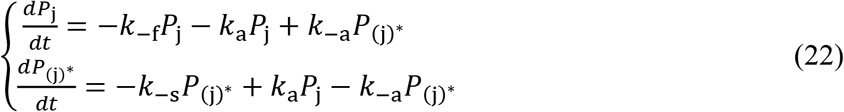

where the subscript j, as above, stands for TCR–pMHC or TCR–pMHC–CD4 bonds in the weak (without *) or strong (with *) state, and all rates *k_i_* vary as a function of force as given by Eq. 21. This two-state catch-bond model could recapitulate the average bond lifetime as well as the biexponentially distributed bond survival probabilities of both TCR–pMHC and TCR–pMHC–CD4 interactions. All kinetic parameters were determined by globally fitting the models to the force-lifetime distributions as a whole via maximum likelihood estimation (MLE). The initial proportions of the two states a *t* = 0 were determined from the ratios of the kinetic rates assuming equilibrium. A two-step optimization procedure was employed to minimize the MLE, which explored a large parameter space by Differential Evolution to obtain an approximate solution, followed by L-BFGS to home in the minimum. This returns the best-fit parameters summarized in Supplementary Fig. 3d. The force dependence of the kinetic rates is graphically depicted in Supplementary Fig. 3c. Using Eqs. 21&22 with the best-fit parameters we calculated the probability of the strong state and its fraction as functions of time and force for both TCR–pMHC and TCR–pMHC–CD4 interactions (Supplementary Fig. 3e-f). The same equations and best-fit parameters predict the mean ± SE of lifetime vs force of the TCR–pMHC, pMHC–CD4 and TCR–pMHC–CD4 bonds that agree well with the mean ± SEM data of these interactions (Fig. 2e). Also predicted are the average lifetime vs force of weak state, strong state, and their sum of the trimolecular bond that shows the force enhancement of cooperativity (Fig. 2f). It is important to note that although the two-state, two-path model fits the data as a catch-slip bond, this does not need to always be the case. In fact, the two-state, two-path model could in theory also yield slip bond or slip-catch-slip bond behavior given specific sets of parameters. Nevertheless, the catch-slip behavior naturally arises from maximizing the likelihood of the TCR–pMHC and TCR–pMHC–CD4 force-lifetime distributions, in good agreement with the average lifetimes of each bin obtained independently from the fitting. This further adds confidence both in the existence of the two catch-bonds as well as the reliability of the model. The standard error for each parameter was obtained by inverting the Hessian matrix of the negative log likelihood at the MLE to obtain the approximate covariance matrix whose diagonal entries correspond to the variance of the fitted parameters. All kinetic modeling and associated statistics were programmatically implemented in Julia and Python.

## Data Availability

Data supporting the findings of this study are presented in the article and supplementary materials and are available from the corresponding authors upon reasonable request.

## Code Availability

The code used to fit the data presented in this study is available at https://github.com/vaglino/TCR-MHC-CD4_kinetic_modeling under MIT license.

## Acknowledgements

We thank the NIH Tetramer Facility at Emory University for providing the pMHC molecules. This work was supported by NIH grants R21AI135753 (to Y.K and C.Z.), R01GM124489, U01CA214354, and U01CA250040 (to C.Z.), and R01AI129893 (to R.A.M.).

## Author Contributions

M.N.R., V.P., C.Z., and Y.K. conceived and designed the study; C.Z. and Y.K. directed the study; M.N.R., V.P., K.L., H.-K.C., J.H., F.G., and P.A. performed experiments; C.Z. and S.T. developed mathematical models; S.T. analyzed data using mathematical model; R.A.M. provided critical reagents; M.N.R., V.P., K.L. Y.K., and C.Z. wrote the paper with contributions from other authors.

## Methods

Fig. 1d: Micropipette adhesion frequency assay with fitting to Eq. 8.

Fig. 2a,b: Micropipette adhesion frequency assay with fitting to Eqs. 1 and 2.

Fig. 2d: BFP thermal fluctuation assay with fitting to ln(*p*) = -*k*_off_ *t*.

Fig. 3a,b,d,e: BFP adhesion frequency assay with the single-species data fitting to Eq. 8 and dual-species data fitting to Eqs. 1 and 2.

Fig. 3e, brown: BFP adhesion frequency assay with the black data fitting to Eqs. 1 and 2 using only two parameters (*K*_a^*^,CD4_ and *k*_−2,CD4_) after fixing *A*_c_*K*_a,TCR_ and *k*_−1,TCR_ at values obtained from fitting the green and purple data by Eq. 8.

Fig 3h: Micropipette adhesion frequency assay with fitting to Eq. 8 (green curve) and Eqs. 1 and 2 (green and black curves, orange curve, and blue curve with *k*_−1,TCR_ and *k*_−2,CD4_ fixed at the values obtained from fitting the orange curve).

Fig. 5a: Micropipette adhesion frequency assay with fitting to Eq. 4. at steady-state.

Fig. S1a-c: Micropipette adhesion frequency assay with fitting to Eq. 7 at steady-state.

Fig. S2c: Micropipette adhesion frequency assay with fitting to Eq. 3. at steady-state.

## References

1 Doyle, C. & Strominger, J. L. Interaction between CD4 and class II MHC molecules mediates cell adhesion. Nature 330, 256–259, doi:10.1038/330256a0 (1987).

2 Norment, A. M., Salter, R. D., Parham, P., Engelhard, V. H. & Littman, D. R. Cell-cell adhesion mediated by CD8 and MHC class I molecules. Nature 336, 79–81, doi:10.1038/336079a0 (1988).

3 Jiang, N. et al. Two-stage cooperative T cell receptor-peptide major histocompatibility complex-CD8 trimolecular interactions amplify antigen discrimination. Immunity 34, 13–23, doi:10.1016/j.immuni.2010.12.017 (2011).

4 Liu, B. et al. 2D TCR-pMHC-CD8 kinetics determines T-cell responses in a self-antigen-specific TCR system. European journal of immunology 44, 239–250, doi:10.1002/eji.201343774 (2014).

5 Hong, J. et al. A TCR mechanotransduction signaling loop induces negative selection in the thymus. Nature immunology 19, 1379–1390, doi:10.1038/s41590-018-0259-z (2018).

6 Stepanek, O. et al. Coreceptor Scanning by the T Cell Receptor Provides a Mechanism for T Cell Tolerance. Cell 159, 333–345, doi:10.1016/j.cell.2014.08.042 (2014).

7 Jonsson, P. et al. Remarkably low affinity of CD4/peptide-major histocompatibility complex class II protein interactions. P Natl Acad Sci USA 113, 5682–5687, doi:10.1073/pnas.1513918113 (2016).

8 Hong, J. et al. Force-regulated In situ TCR-peptide-bound MHC class II kinetics determine functions of CD4+ T cells. Journal of immunology 195, 3557–3564, doi:10.4049/jimmunol.1501407 (2015).

9 Mørch, A. M., Bálint, Š., Santos, A. M., Davis, S. J. & Dustin, M. L. Coreceptors and TCR Signaling – the Strong and the Weak of It. Frontiers in Cell and Developmental Biology 8, doi:10.3389/fcell.2020.597627 (2020).

10 Xiong, Y., Kern, P., Chang, H. & Reinherz, E. T Cell Receptor Binding to a pMHCII Ligand Is Kinetically Distinct from and Independent of CD4. The Journal of biological chemistry 276, 5659–5667, doi:10.1074/jbc.M009580200 (2001).

11 Liu, B., Chen, W., Evavold, B. D. & Zhu, C. Accumulation of dynamic catch bonds between TCR and agonist peptide-MHC triggers T cell signaling. Cell 157, 357–368, doi:10.1016/j.cell.2014.02.053 (2014).

12 Yin, Y., Wang, X. X. & Mariuzza, R. A. Crystal structure of a complete ternary complex of T-cell receptor, peptide-MHC, and CD4. Proceedings of the National Academy of Sciences 109, 5405–5410, doi:10.1073/pnas.1118801109 (2012).

13 Courtney, A. H., Lo, W. L. & Weiss, A. TCR Signaling: Mechanisms of Initiation and Propagation. Trends in biochemical sciences, doi:10.1016/j.tibs.2017.11.008 (2017).

14 Rudd, C. E. CD4, CD8 and the TCR-CD3 complex: a novel class of protein-tyrosine kinase receptor. Immunology Today 11, 400–406, doi:https://doi.org/10.1016/0167-5699(90)90159-7 (1990).

15 Xu, H. & Littman, D. R. A kinase-independent function of Lck in potentiating antigen-specific T cell activation. Cell 74, 633–643, doi:10.1016/0092-8674(93)90511-n (1993).

16 Glatzova, D. & Cebecauer, M. Dual Role of CD4 in Peripheral T Lymphocytes. Frontiers in immunology 10, 618, doi:10.3389/fimmu.2019.00618 (2019).

17 Wang, X. X. et al. Affinity maturation of human CD4 by yeast surface display and crystal structure of a CD4-HLA-DR1 complex. P Natl Acad Sci USA 108, 15960–15965, doi:10.1073/pnas.1109438108 (2011).

18 Huppa, J. B. et al. TCR-peptide-MHC interactions in situ show accelerated kinetics and increased affinity. Nature 463, 963–967, doi:10.1038/nature08746 (2010).

19 Dong et al. Structural basis of assembly of the human T cell receptor-CD3 complex. Nature, doi:10.1038/s41586-019-1537-0 (2019).

20 Casas, J. et al. Ligand-engaged TCR is triggered by Lck not associated with CD8 coreceptor. Nature communications 5, 5624, doi:10.1038/ncomms6624 (2014).

21 Crawford, F., Kozono, H., White, J., Marrack, P. & Kappler, J. Detection of Antigen-Specific T Cells with Multivalent Soluble Class II MHC Covalent Peptide Complexes. Immunity 8, 675–682, doi:10.1016/s1074-7613(00)80572-5 (1998).

22 Glassman, C. R., Parrish, H. L., Lee, M. S. & Kuhns, M. S. Reciprocal TCR-CD3 and CD4 Engagement of a Nucleating pMHCII Stabilizes a Functional Receptor Macrocomplex. Cell Reports 22, 1263–1275, doi:10.1016/j.celrep.2017.12.104 (2018).

23 Chesla, S. E., Selvaraj, P. & Zhu, C. Measuring Two-Dimensional Receptor-Ligand Binding Kinetics by Micropipette. Biophys. J. 75, 1553–1572 (1998).

24 Huang, J. et al. The kinetics of two-dimensional TCR and pMHC interactions determine T-cell responsiveness. Nature 464, 932–U156, doi:10.1038/nature08944 (2010).

25 Deng, L. et al. Structural basis for the recognition of mutant self by a tumor-specific, MHC class II-restricted T cell receptor. Nat. Immunol. 8, 398–408, doi:10.1038/ni1447 (2007).

26 Huang, J., Edwards, L. J., Evavold, B. D. & Zhu, C. Kinetics of MHC-CD8 interaction at the T cell membrane. J. Immunol. 179, 7653–7662 (2007).

27 Zhu, C. & Williams, T. E. Modeling concurrent binding of multiple molecular species in cell adhesion. Biophysical journal 79, 1850–1857, doi:10.1016/S0006-3495(00)76434-4 (2000).

28 Williams, T. E., Selvaraj, P. & Zhu, C. Concurrent binding to multiple ligands: kinetic rates of CD16b for membrane-bound IgG1 and IgG2. Biophysical journal 79, 1858–1866, doi:10.1016/S0006-3495(00)76435-6 (2000).

29 Williams, T. E., Nagarajan, S., Selvaraj, P. & Zhu, C. Concurrent and independent binding of Fcgamma receptors IIa and IIIb to surface-bound IgG. Biophysical journal 79, 1867–1875, doi:10.1016/S0006-3495(00)76436-8 (2000).

30 Chen, W., Evans, E. A., McEver, R. P. & Zhu, C. Monitoring receptor-ligand interactions between surfaces by thermal fluctuations. Biophys. J. 94, 694–701 (2008).

31 Marshall, B. T. et al. Direct observation of catch bonds involving cell-adhesion molecules. Nature 423, 190–193, doi:10.1038/nature01605 (2003).

32 Kao, H. & Allen, P. M. An antagonist peptide mediates positive selection and CD4 lineage commitment of MHC class II-restricted T cells in the absence of CD4. J. Exp. Med. 201, 149–158, doi:10.1084/jem.20041574 (2005).

33 Rothemund, P. W. Folding DNA to create nanoscale shapes and patterns. Nature 440, 297–302, doi:10.1038/nature04586 (2006).

34 Williams, T. E., Nagarajan, S., Selvaraj, P. & Zhu, C. Quantifying the impact of membrane microtopology on effective two-dimensional affinity. The Journal of biological chemistry 276, 13283–13288, doi:10.1074/jbc.M010427200 (2001).

35 Li, Q.-J. et al. CD4 enhances T cell sensitivity to antigen by coordinating Lck accumulation at the immunological synapse. Nat Immunol 5, 791–799 (2004).

36 Natarajan, K. et al. An allosteric site in the T-cell receptor Cbeta domain plays a critical signalling role. Nature communications 8, 15260, doi:10.1038/ncomms15260 (2017).

37 He, Y. et al. Peptide-MHC Binding Reveals Conserved Allosteric Sites in MHC Class I-and Class II-Restricted T Cell Receptors (TCRs). J Mol Biol 432, 166697, doi:10.1016/j.jmb.2020.10.031 (2020).

38 Rangarajan, S. et al. Peptide-MHC (pMHC) binding to a human antiviral T cell receptor induces long-range allosteric communication between pMHC-and CD3-binding sites. The Journal of biological chemistry 293, 15991–16005, doi:10.1074/jbc.RA118.003832 (2018).

39 Zhu, C., Chen, W., Lou, J., Rittase, W. & Li, K. Mechanosensing through immunoreceptors. Nature immunology, doi:10.1038/s41590-019-0491-1 (2019).

40 Li, K. et al. PD-1 suppresses TCR-CD8 cooperativity during T-cell antigen recognition. Nature communications 12, 2746, doi:10.1038/s41467-021-22965-9 (2021).

41 Rudd, C. E., Trevillyan, J. M., Dasgupta, J. D., Wong, L. L. & Schlossman, S. F. The CD4 receptor is complexed in detergent lysates to a protein-tyrosine kinase (pp58) from human T lymphocytes. Proc Natl Acad Sci U S A 85, 5190–5194, doi:10.1073/pnas.85.14.5190 (1988).

42 Burgess, K. E. et al. Biochemical identification of a direct physical interaction between the CD4:p56lck and Ti(TcR)/CD3 complexes. European journal of immunology 21, 1663–1668, doi:10.1002/eji.1830210712 (1991).

43 Duplay, P., Thome, M., Herve, F. & Acuto, O. P56(Lck) Interacts Via Its Src Homology-2 Domain with the Zap-70 Kinase. Journal of Experimental Medicine 179, 1163–1172, doi:DOI 10.1084/jem.179.4.1163 (1994).

44 Thome, M., Germain, V., DiSanto, J. P. & Acuto, O. The p56lck SH2 domain mediates recruitment of CD8/p56lck to the activated T cell receptor/CD3/zeta complex. European journal of immunology 26, 2093–2100, doi:10.1002/eji.1830260920 (1996).

45 Chen, W., Zarnitsyna, V. I., Sarangapani, K. K., Huang, J. & Zhu, C. Measuring Receptor-Ligand Binding Kinetics on Cell Surfaces: From Adhesion Frequency to Thermal Fluctuation Methods. Cell Mol Bioeng 1, 276–288, doi:10.1007/s12195-008-0024-8 (2008).

46 Evans, E., Leung, A., Heinrich, V. & Zhu, C. Mechanical switching and coupling between two dissociation pathways in a P-selectin adhesion bond. Proceedings of the National Academy of Sciences of the United States of America 101, 11281–11286, doi:10.1073/pnas.0401870101 (2004).

47 Buckley, C. D. et al. Cell adhesion. The minimal cadherin-catenin complex binds to actin filaments under force. Science 346, 1254211, doi:10.1126/science.1254211 (2014).

48 Bell, G. Models for the specific adhesion of cells to cells. Science 200, 618–627, doi:10.1126/science.347575 (1978).

