## Supplemental Info for "Cooperative binding of TCR and CD4 to pMHC enhances TCR sensitivity"

Supplementary Figure 1

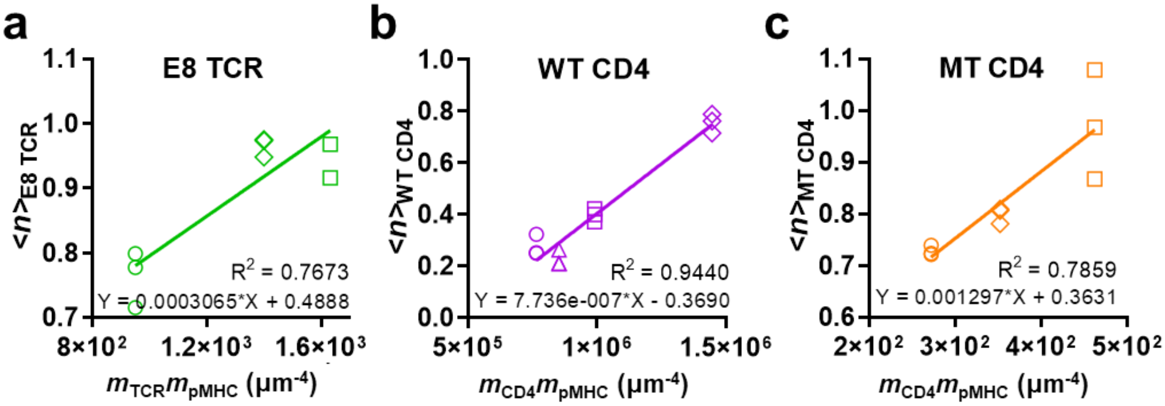


**Supplementary Figure 1 | Average number of bonds are proportional to receptor and ligand densities. Related to Fig. 1.** Average number of bonds at steady-state vs the product of receptor and ligand densities are fitted by linear regression for pMHC interaction with E8 TCR (**a**), WT CD4 (**b**), or MT CD4 (**c**). The slopes, representing $A_cK_{a,TCR}$, $A_cK_{a,CD4}$, and $A_cK_{a,MT\ CD4}$, respectively, are listed in Table 1.

### Supplementary Figure 2

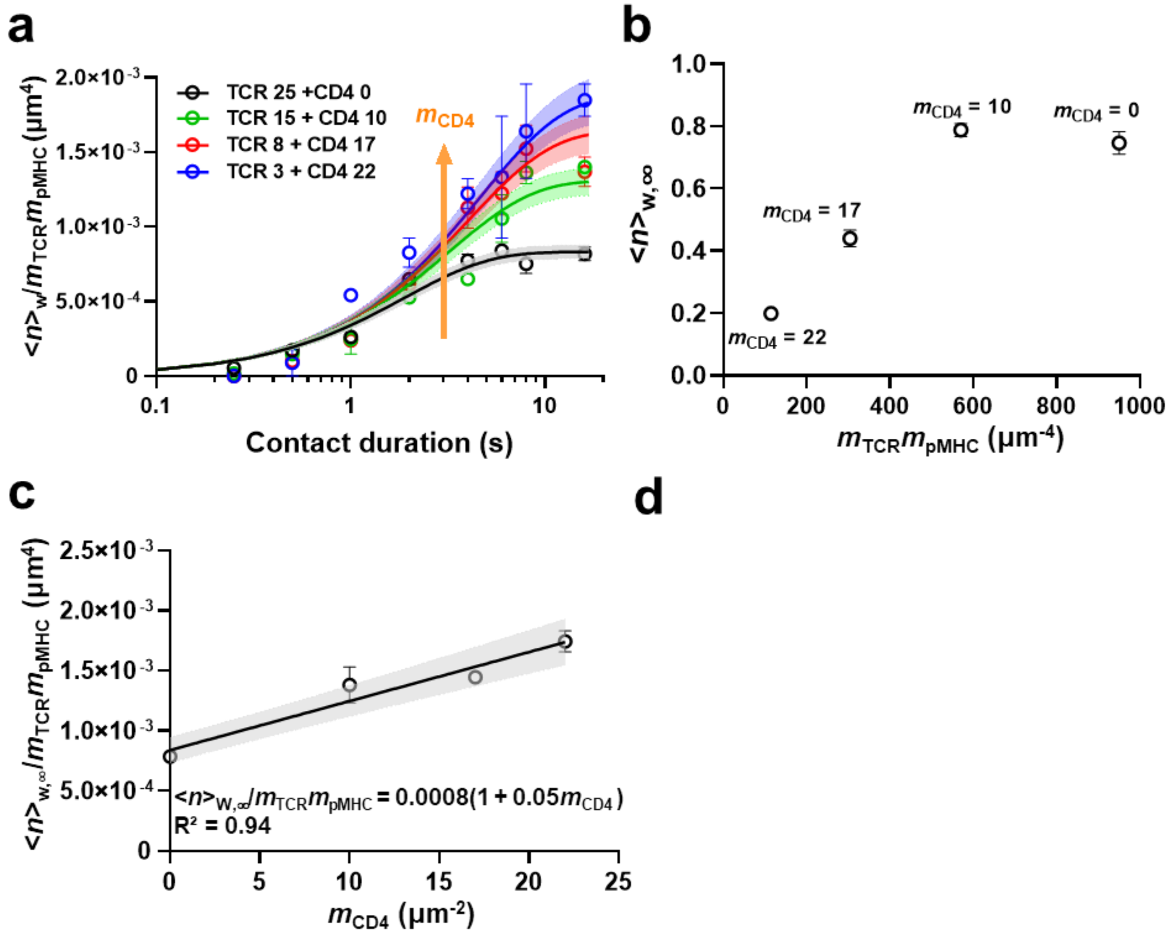


**Supplementary Figure 2 | TCR-CD4 cooperativity analysis. Related to Fig. 2b. a,** The same data in Figure 2b are plot on semi-log scale. **b,** Average bond number at steady-state vs the product of TCR and pMHC densities for data in Figure 2a. **c,** Normalized whole number of bonds at steady-state is proportional to CD4 density. According to Eq. 3, slope and *y*-intercept represent $K^{*}_{a,CD4}$ and $A_c K_{a,TCR}$, respectively, and are listed in Table 1.

### Supplementary Figure 3

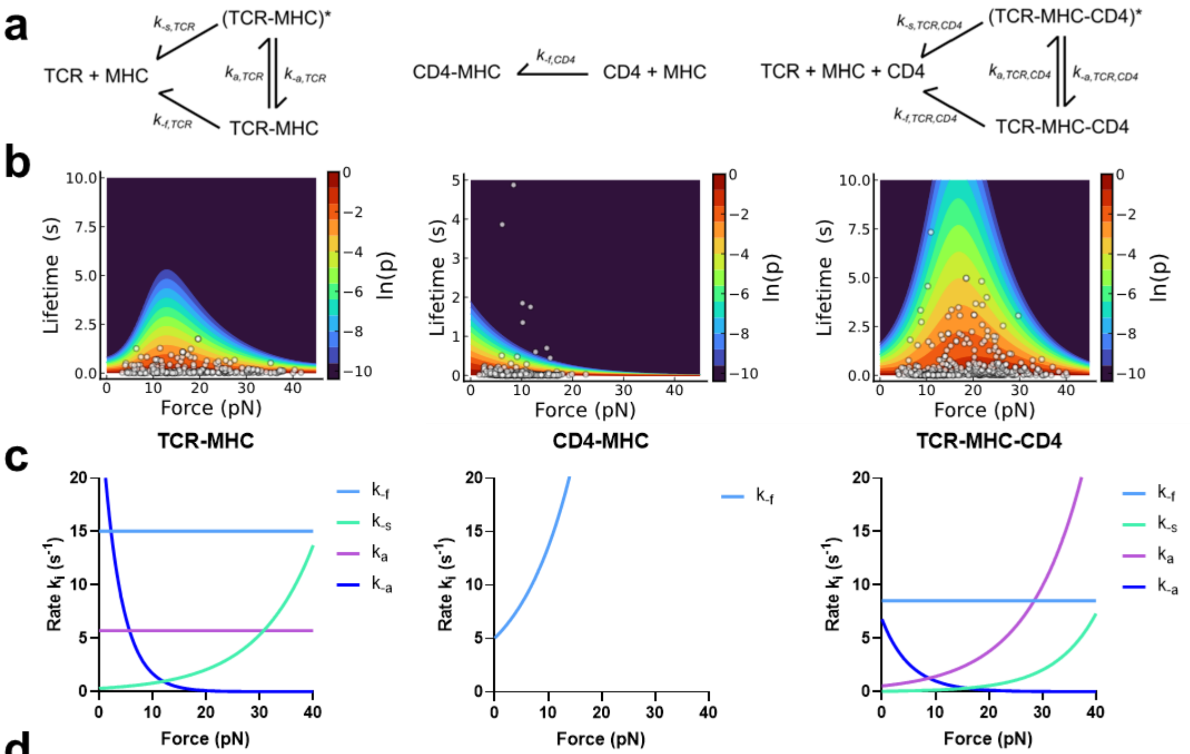


d

| Transition | Rate | $k^0$ (1/s) | $x^0$ (Å) |
|---|---|---|---|
| TCR–MHC → 0 | $k_{f,TCR}$ | 15.0 ± 0.616 | 0 ± 0.0644 |
| (TCR–MHC)* → 0 | $k_{s,TCR}$ | 0.450 ± 0.0638 | 3.39 ± 0.235 |
| TCR–MHC → (TCR–MHC)* | $k_{a,TCR}$ | 6.37 ± 1.19 | 0 ± 0.545 |
| TCR–MHC ← (TCR–MHC)* | $k_{-a,TCR}$ | 31.1 ± 6.30 | -13.1 ± 1.28 |

| Transition | Rate | $k^0$ (1/s) | $x^0$ (Å) |
|---|---|---|---|
| CD4–MHC → 0 | $k_{f,CD4}$ | 5.03 ± 0.427 | 4.08 ± 0.387 |

| Transition | Rate | $k^0$ (1/s) | $x^0$ (Å) |
|---|---|---|---|
| TCR–MHC–CD4 → 0 | $k_{f,TCR,CD4}$ | 8.53 ± 0.190 | 0 ± 0.0556 |
| (TCR–MHC–CD4)* → 0 | $k_{s,TCR,CD4}$ | 0.0450 ± 0.00769 | 5.24 ± 0.223 |
| TCR–MHC–CD4 → (TCR–MHC–CD4)* | $k_{a,TCR,CD4}$ | 0.544 ± 0.0578 | 3.98 ± 0.219 |
| TCR–MHC–CD4 ← (TCR–MHC–CD4)* | $k_{-a,TCR,CD4}$ | 6.80 ± 0.743 | -7.83 ± 0.528 |

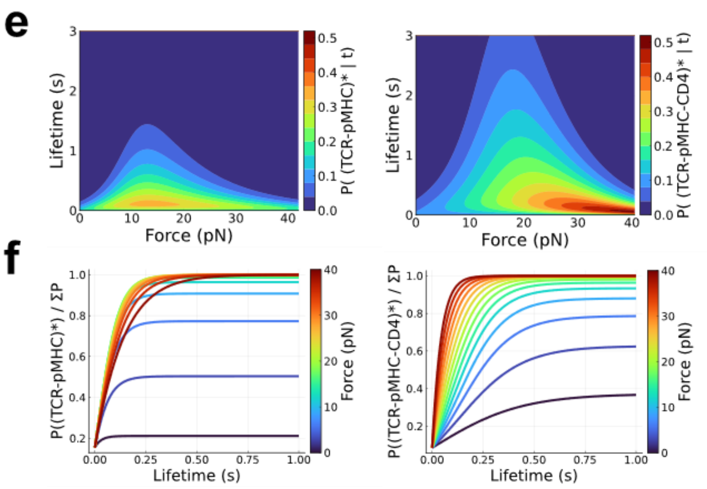

**Supplementary Figure 3 | Two-state catch bond models fit the force-dependent lifetime data from experiments. Related to Fig. 2d-f. a,** Schematics of two two-state catch bond models and a single-state slip bond model with indicated reaction rates for TCR–pMHC (*left*) and CD4–pMHC (*middle*) bimolecular dissociations, and for the case of TCR–pMHC–CD4 trimolecular dissociation (*right*). The superscript * labels the strong states. **b,** Bond lifetime vs force scattergrams of TCR–pMHC (*left*), CD4–pMHC (*middle*), and TCRαβ + CD4 vs pMHC (*right*) interactions fitted using maximum likelihood estimation (MLE). Depending on the Bell parameters, the three kinetic models generate three families of force-lifetime curves. The likelihoods for data (points) to be observed as predicted (curves) were maximized to obtain the best-fit parameters. Parameters so evaluated from global MLE fitting predict mean ± SE curves of bond lifetime vs force shown in Fig. 2e. They also predict the lifetimes vs force of TCR–pMHC–CD4 trimolecular bonds in the strong state, weak state, and their sum as shown in Fig. 2f. **c,** Reaction rates vs force plots. The force-dependent rates for fast and slow dissociations as well as for activation and deactivation state transitions are plotted for the TCR–pMHC (*left*), CD4–pMHC (*middle*), and TCR–pMHC–CD4 (*right*) bonds. The plots are shown with the best-fit parameters. **d**, Summary of the best-fit Bell parameters used to model the TCR–pMHC (*top*), CD4–pMHC (*middle*), TCR–pMHC–CD4 (*bottom*) dissociations. **e, f**, Probability (e) and fraction (f) of strong bonds as a function of time and force for TCR–pMHC (*left*) and TCR–pMHC–CD4 (*right*) bonds.

### Supplementary Figure 4

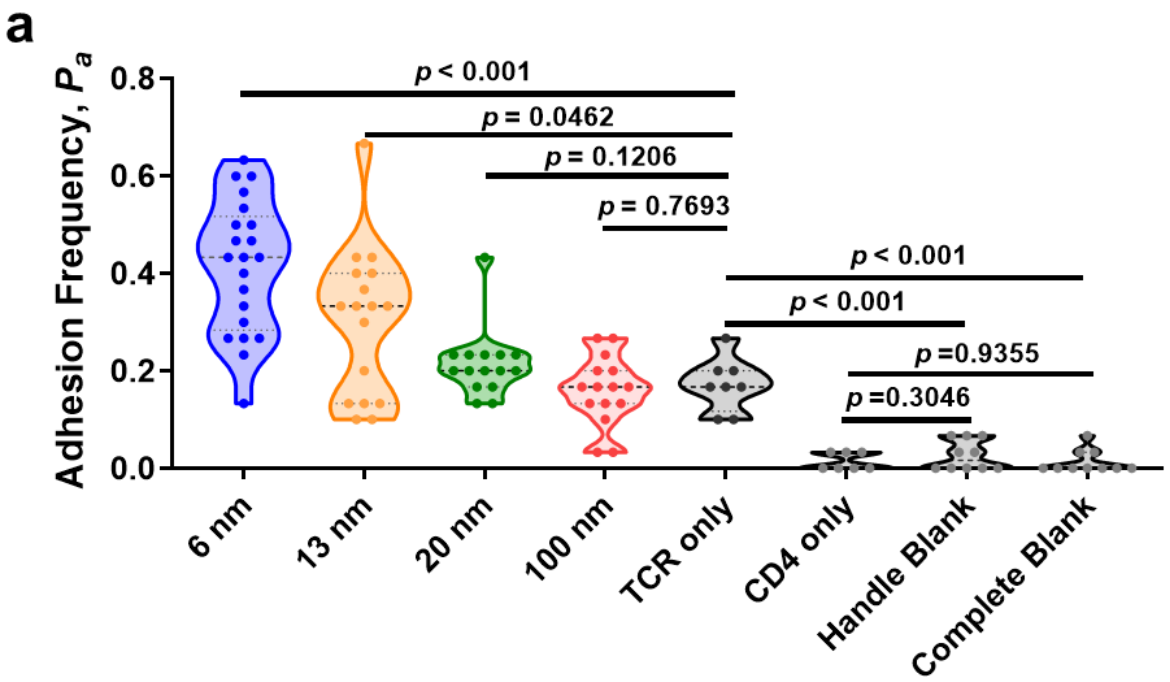


**Supplementary Figure 4 | TCR-CD4 cooperativity analysis using DNA origami. Related to Fig. 4.** Adhesion frequencies ($P_a$) of RBCs bearing pMHC interacting with DNA origami beads presenting TCR and CD4 at 6, 13, 20, or 100 nm spacing, or presenting TCR or CD4 alone, or no protein (handle blank and complete blank) at contact time of 4 s (n = 21, 17, 15, 15, 8, 7, 10, and 10 cell-bead pairs). In the violin plots that show data densities, the dashed lines in the middle represent mean and the lower and upper dotted lines represent the first and third quartiles. P-values were calculated for indicated groups using the Mann-Whitney test.

### Supplementary Table 1 | Site densities of pMHC, TCR, and CD4.

| | $m_{TCR}$ (#/μm$^2$) | $m_{CD4}$ (#/μm$^2$) | $m_{pMHC}$ (#/μm$^2$) |
|---|---|---|---|
| Fig. 1b orange | 0 | 16 | 17 |
| Fig. 1b green | 25 | n/a | 38 |
| Fig. 1b purple | 0 | 900 | 850 |
| Fig. 1c orange | 0 | 16 | 17 |
| Fig. 1c green | 0 | 22 | 16 |
| Fig. 1c purple | 0 | 21 | 22 |
| Fig. 1d orange square | 0 | 16 | 22 |
| Fig. 1d orange triangle | 0 | 16 | 17 |
| Fig. 1d orange circle | 0 | 21 | 22 |
| Fig. 1d green square | 50 | 0 | 28 |
| Fig. 1d green triangle | 48 | 0 | 34 |
| Fig. 1d green circle | 25 | 0 | 38 |
| Fig. 1d purple square | 0 | 900 | 1100 |
| Fig. 1d purple triangle | 0 | 900 | 850 |
| Fig. 1d purple circle | 0 | 1700 | 850 |
| Fig. 1d purple diamond | 0 | 1550 | 550 |
| Fig. 2a black | 25 | 0 | 38 |
| Fig. 2a green | 15 | 10 | 38 |
| Fig. 2a red | 8 | 17 | 38 |
| Fig. 2a blue | 3 | 22 | 38 |
| Fig. 3a green | 58 | 0 | 19 |
| Fig. 3a purple | 0 | 1948 | 19 |
| Fig. 3a black | 76 | 96 | 8 |
| Fig. 3c green | 58 | 0 | 19 |
| Fig. 3c purple | 0 | 1948 | 19 |
| Fig. 3c black | 76 | 96 | 8 |
| Fig. 3d black | 14 | 5 | 207 |
| Fig. 3d green | 25 | 0 | 200 |
| Fig. 3d purple | 0 | 10 | 2832 |
| Fig. 3f green | 35 | 0 | n/a |
| Fig. 3f purple | 0 | 10 | n/a |
| Fig. 3f black | 7 | 7 | n/a |
| Fig. 3f blue | 35 | 9 | n/a |
| Fig. 3h black | 14 | 11 | 29 |
| Fig. 3h green | 42 | 0 | 5 |
| Fig. 3h orange | 49 | 53 | 4 |
| Fig. 3h blue | 2 | 2 | 52 |
| Fig. 5a blue | 35 | 35 | 460 |
| Fig. 5a orange | 32 | 32 | 530 |
| Fig. 5a green | 29 | 29 | 530 |
| Fig. 5a red | 30 | 30 | 550 |
| Fig. 5a black-TCR | 23 | n/a | 563 |
| Fig. 5a black-CD4 | n/a | 23 | 563 |
